# Paleoclimate and current climate collectively shape the phylogenetic and functional diversity of trees worldwide

**DOI:** 10.1101/2020.06.02.128975

**Authors:** Wen-Yong Guo, Josep M. Serra-Diaz, Franziska Schrodt, Wolf L. Eiserhardt, Brian S. Maitner, Cory Merow, Cyrille Violle, Anne Blach-Overgaard, Jian Zhang, Madhur Anand, Michaël Belluau, Hans Henrik Bruun, Chaeho Byun, Jane A. Catford, Bruno E. L. Cerabolini, Eduardo Chacón-Madrigal, Daniela Ciccarelli, Johannes H. C. Cornelissen, Anh Tuan Dang-Le, Angel de Frutos, Arildo S. Dias, Aelton B. Giroldo, Alvaro G. Gutiérrez, Wesley Hattingh, Tianhua He, Peter Hietz, Nate Hough-Snee, Steven Jansen, Jens Kattge, Tamir Klein, Benjamin Komac, Nathan Kraft, Koen Kramer, Sandra Lavorel, Christopher H. Lusk, Adam R. Martin, Maurizio Mencuccini, Sean T. Michaletz, Vanessa Minden, Akira S. Mori, Ülo Niinemets, Yusuke Onoda, Renske E. Onstein, Josep Peñuelas, Valério D. Pillar, Jan Pisek, Matthew J. Pound, Bjorn J.M. Robroek, Brandon Schamp, Martijn Slot, Ênio Sosinski, Nadejda A. Soudzilovskaia, Nelson Thiffault, Peter van Bodegom, Fons van der Plas, Ian J. Wright, Jingming Zheng, Brian J. Enquist, Jens-Christian Svenning

## Abstract

Trees are of vital importance for ecosystem functioning and services at local to global scales, yet we still lack a detailed overview of the global patterns of tree diversity and the underlying drivers, particularly the imprint of paleoclimate. Here, we present the high-resolution (110 km) worldwide mapping of tree species richness, functional and phylogenetic diversities based on ∼7 million quality-assessed occurrences for 46,752 tree species (80.5% of the estimated total number of tree species), and subsequent assessments of the influence of paleo-climate legacies on these patterns. All three tree diversity dimensions exhibited the expected latitudinal decline. Contemporary climate emerged as the strongest driver of all diversity patterns, with Pleistocene and deeper-time (>10^7^ years) paleoclimate as important co-determinants, and, notably, with past cold and drought stress being linked to reduced current diversity. These findings demonstrate that tree diversity is affected by paleoclimate millions of years back in time and highlight the potential for tree diversity losses from future climate change.

---

Understanding the global distribution of tree diversity and its underlying drivers has been an enduring pursuit of scientists, at least as far back as Alexander von Humboldt. Achieving this aim is becoming ever more urgent due to forest degradation and land use change ^1–4^, and also for aiding forest restoration efforts ^5^. Although our understanding of the global extent of tree cover has been greatly improved via remote sensing ^6^ and large networks of forest tree plots ^7,8^, there are still gaps in our knowledge of the global patterns and drivers of tree diversity. Previous studies have often focused on tree species richness (SR) (e.g., ^9^). However, SR does not directly represent species’ evolutionary history and does not provide trait-based insight into their functioning and role in ecosystems. Phylogenetic diversity (PD; ^10,11^) and functional diversity (FD; ^12^) have been introduced as promising, more informative biodiversity variables than SR and have been successfully used in a wide variety of ecological applications, including conservation prioritization ^13–15^. Indeed, FD is better coupled than SR to ecosystem functioning, e.g., productivity responses to climate change and forest multi-functionality ^16–18^. In addition, PD and FD are more informative than SR in describing mechanisms of species coexistence and ecosystem functioning ^19,20^, and thus shed light on species extinction and conservation ^13,14,21–23^.

Many studies have emphasized the importance of current climate and edaphic conditions as key determinants of species diversity (e.g., ^24,25^). However, paleoclimate could leave an influential legacy, e.g. via speciation, extinction or dispersal, on contemporary SR, and on phylogenetic and functional structures of forest ecosystems ^8,26–32^. Earth climate has experienced continuous changes during geological time ^33^, such as cooling or warming trends and events, and major climatic transitional periods have coincided with global ecosystem shifts ^34–37^. Climatically-stable regions tend to have high speciation and low extinction rates, resulting in higher SR, FD, and PD ^38,39^. Contrastingly, wide climate oscillations (like glacial-interglacial cycles) can dramatically truncate species’ ranges and the chances of local diversification and adaptation, increasing the likelihood of extinction and the removal of species with suboptimal traits, thereby decreasing all three facets of diversity ^40–43^. However, rapid climate change may alternatively cause range fragmentation and further allopatric speciation as the result of isolation, potentially increasing net diversification rates ^27^.

Due to the non-equivalency between the facets of diversity ^44–47^, the responses of SR, PD and FD to different climatic conditions may vary. For example, warm and humid climates are hypothesized to increase diversification rates ^48,49^, dispersal and establishment ^50^, and decrease extinction ^27,51^, thus increasing SR and PD, but not necessarily FD, as comparable climates more likely predispose species towards similar functional traits ^25,52–55^. Thus, contemporary species diversity patterns can be the result of historical climate legacies and present-day environment, although the relative importance of these factors for FD and PD could be different.

Variable geological climates, i.e., warm and humid, or cold and dry in different paleo-time periods, had remarkably divergent influences on tree diversities. However, previous studies have concentrated mostly on assessing the effect of the cold and dry Last Glacial Maximum (LGM) imprints that occurred ∼27 – 19 thousand years ago (kya), but deeper-time perspectives may also be important. For instance, ref. ^26^ found that palm tree diversity in Africa was affected by deep-time climate during the late Pliocene (3.3 – 3.0 million years ago [mya]) and the late Miocene (11.6 – 7.3 mya), respectively. Similarly, ref. ^27^ found that the late Miocene climate influenced global patterns of conifer phylogenetic structure. Recently, ref. ^31^ reported opposite effects of LGM and Miocene tree cover on tree phylogenetic endemism. Hence, considering paleoclimate jointly across a range of time frames could be helpful in better understanding the factors shaping tree diversities. However, only a few SR studies have explicitly considered this ^26^, and even fewer in FD and PD research ^42^.

Here, we go beyond global mapping of tree species richness ^9^ by estimating species composition and, based thereon, functional and phylogenetic diversity. We subsequently analyze the relative roles of past and present climates in shaping global patterns of tree SR, FD, and PD. We first compiled the most updated dataset of tree species including occurrence records, functional traits, and tree phylogeny, covering 46,752 tree species or 80.5% of the species in the GlobalTreeSearch list ^47,56,57^. We subsequently mapped global tree SR, FD, and PD. To understand the potential effects of paleoclimatic change on tree diversities completely, we examined the relative importance of three paleoclimatic states in determining current SR, FD and PD patterns, with consideration of other potential contemporary covariates, such as current climate, elevation, and human activities (Table S1). Specifically, we explored the influence of paleoclimate related to important climate states of the late Cenozoic, the time frame where current species diversity to a large extent have evolved: i) the warm and humid late Miocene, *ca*. 11.63 – 7.25 mya; the mid-Pliocene Warm period, *ca*. 3.264 – 3.025 mya; the cold and dry Pleistocene glaciations (represented by the LGM, ∼ 21 kya); and Pleistocene warm interglacials (IG, ∼ 787 kya and ∼ 130 kya) (Figs. S1 & S2). In doing so, our study addressed three main goals: (1) mapping global contemporary tree SR, FD and PD; (2) assessing the relative importance of present-day environment, Quaternary glacial-interglacial oscillations, and deeper-time effects on today’s SR, FD and PD patterns, to help understand the fundamental processes determining accumulation and maintenance of tree diversity; and (3) investigating spatial divergence between FD and PD, and identifying the underlying driving factors.

## Results

### Global patterns of tree diversities

The global tree SR, FD, and PD distributions show classic latitudinal gradients ^58–60^, with low diversities at high latitudes and the highest diversities in the tropics (grid cell maximum value of 3261 spp. for SR and cumulative branch lengths of 641 and 61,183 Myrs for FD and PD, respectively at 110 km resolution, Fig. S3), particularly in the Neotropical lowlands (Amazonia) (Fig. 1). The latitudinal pattern is stronger in America and Asia-Australasia than in Africa-Europe, due to the interruption of the diversity gradients by deserts in northern Africa, where the diversity indices (SR, FD, and PD) are as low as at latitudes harboring the boreal climate. The similarity of the spatial patterns among the three diversity measures reflects the monotonic relationships observed between them (Fig. S4).

**Fig. 1.**
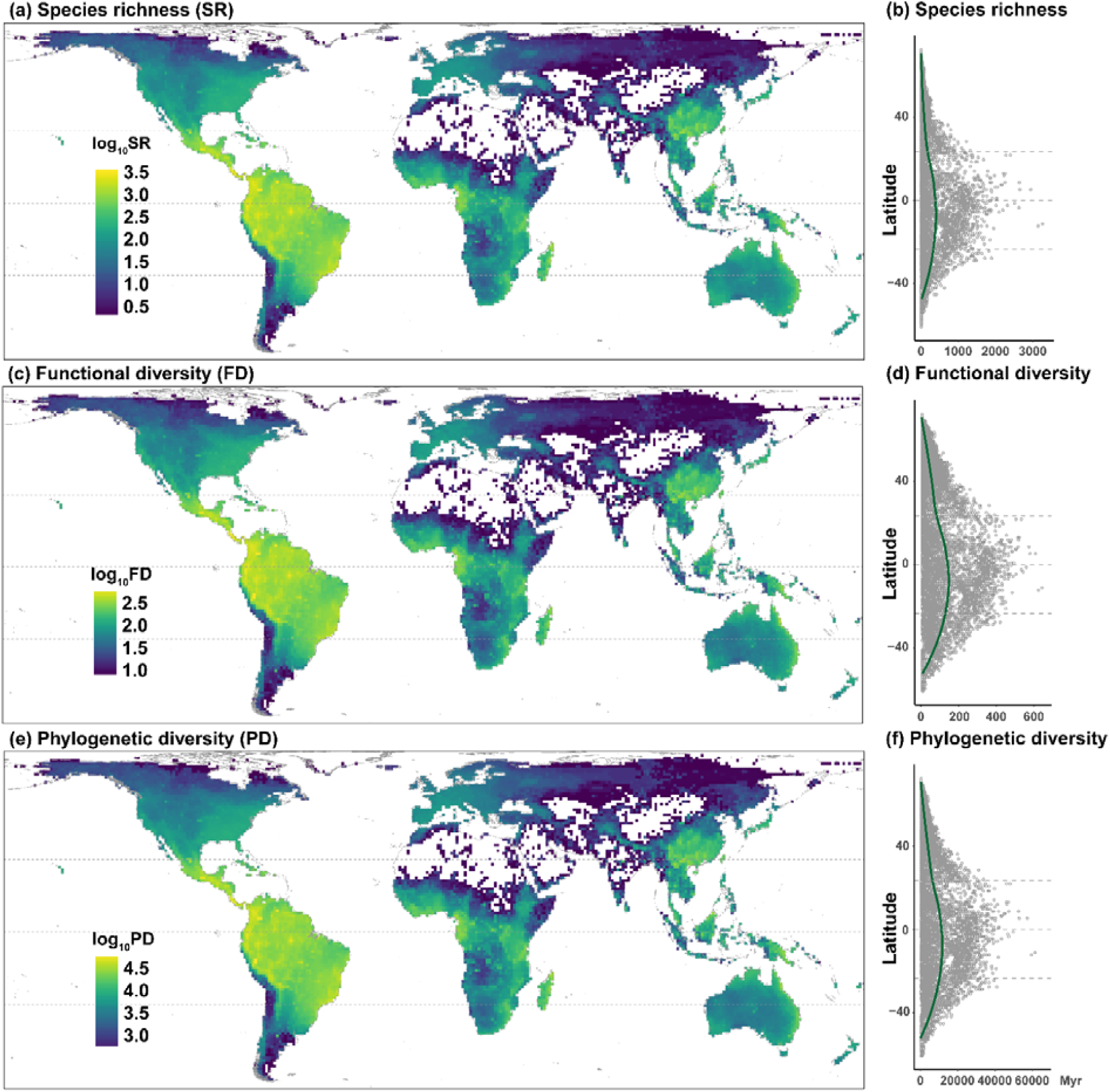
Global patterns of tree (a) species richness, (c) functional diversity, and (e) phylogenetic diversity. In (b), (d) and (f), the fitted line is the lowess regression. Maps use the Behrmann projection at 110 km × 110 km spatial resolution. Myr: Million years.

### Drivers of global tree diversity

Due to the high associations between SR, FD, and PD, their individual relationships with the tested predictors are mostly consistent (Figs. 2 & Table S2). After controlling for spatial autocorrelation, simultaneous autoregressive models (SARs) explain more than 94% (global models) and 78% (regional models) of the variance (Table S2) in the response variables (SR, PD, and FD). Present-day annual precipitation (AP) and mean annual temperature (MAT) are the overall strongest drivers with positive effects on SR, FD, and PD globally, and for AP also regionally except for two regions where other drivers are stronger (Australasia, Nearctic). The effect of MAT varies in strength and sign among regions, showing both positive and negative effects on diversity (Fig. 2, Table S2). Elevation range and human modification index (HMc) have consistent positive effects on SR, FD and PD globally as well as regionally. Four out of the six paleoclimatic variables show significant relations to all three diversity dimensions (Fig. 2). Globally, the Miocene MAT anomaly (i.e., Miocene MAT minus present MAT), the Miocene AP anomaly, and the LGM AP anomaly have positive relations to all diversity indices, while the LGM MAT anomaly have a weak negative relation to SR (*p* < 0.05, Table S2) and no relation to FD and PD (Fig. 2). Hence, SR, PD and FD consistently increase with increasing high precipitation in the Miocene and LGM relative to the present, while SR, but not FD or PD, is generally reduced by increasing warm during LGM at a global scale. However, although some of these global relationships are mirrored regionally, not all paleoclimatic predictors are significant nor show consistent relationships across the biogeographic regions, e.g., with LGM AP anomaly showing negative associations in Australasia and Miocene MAT anomaly in Afrotropic for all three indices (Table S2).

**Fig. 2.**
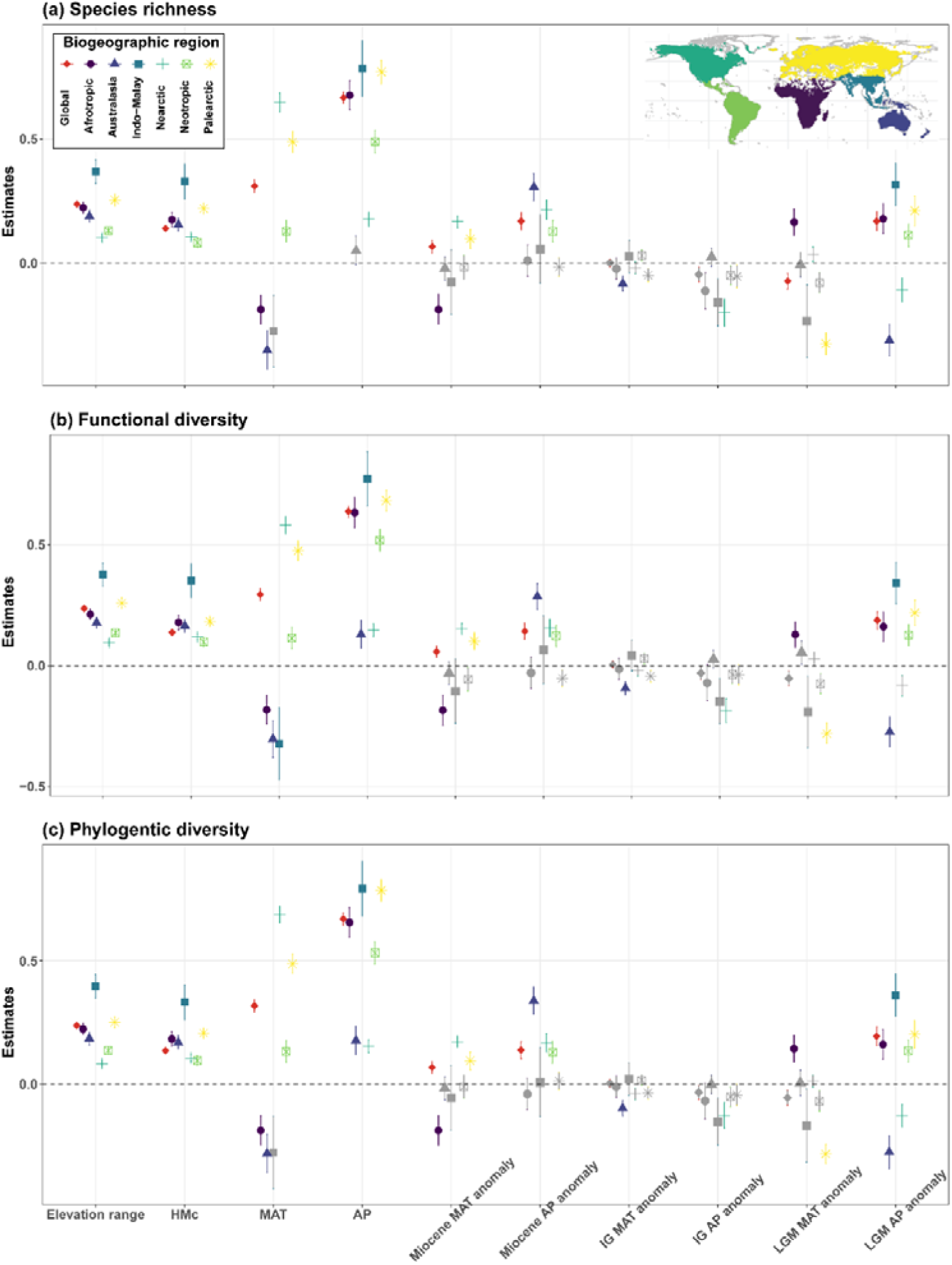
Effects of the tested environmental variables on tree (a) specie richness (SR), (b) functional diversity (FD) and (c) phylogenetic diversity (PD). Estimates (± 1standard error) of effects were obtained from simultaneous autoregressive (SAR) models. Different colors and shapes indicate biogeographic regions. Non-significant variables (*p* > 0.05) are indicated in grey. Results from OLS models are shown in Table S2. HMc: human modification index; MAT: mean annual temperature; AP: Annual precipitation; IG: Pleistocene Interglacial; LGM: Last Glacial Maximum. Anomaly was calculated as the past minus the present state.

Taken all together, precipitation-related effects were stronger and more consistent (among regions) climatic drivers of diversity (SR, FD, PD) than were temperature-related effects, with this true both for current climate (AP) and for paleoclimates (Miocene AP anomaly; IG AP anomaly).

### Spatial divergence between functional and phylogenetic diversities and its drivers

FD and PD are tightly and positively related (Fig. S4c). Deviations (FD residuals) from this linear relationship show marked spatial patterning (Fig. 3). Across North America, western and southern Europe, central Africa, eastern Asia, and eastern Australia, FD is generally higher than predicted by PD (i.e., overdispersion), whilst the opposite (i.e., FD deficit) is revealed in western Australia, much of southern and eastern Africa, west of the Andes (Peru), and central parts of northern Eurasia.

**Fig. 3.**
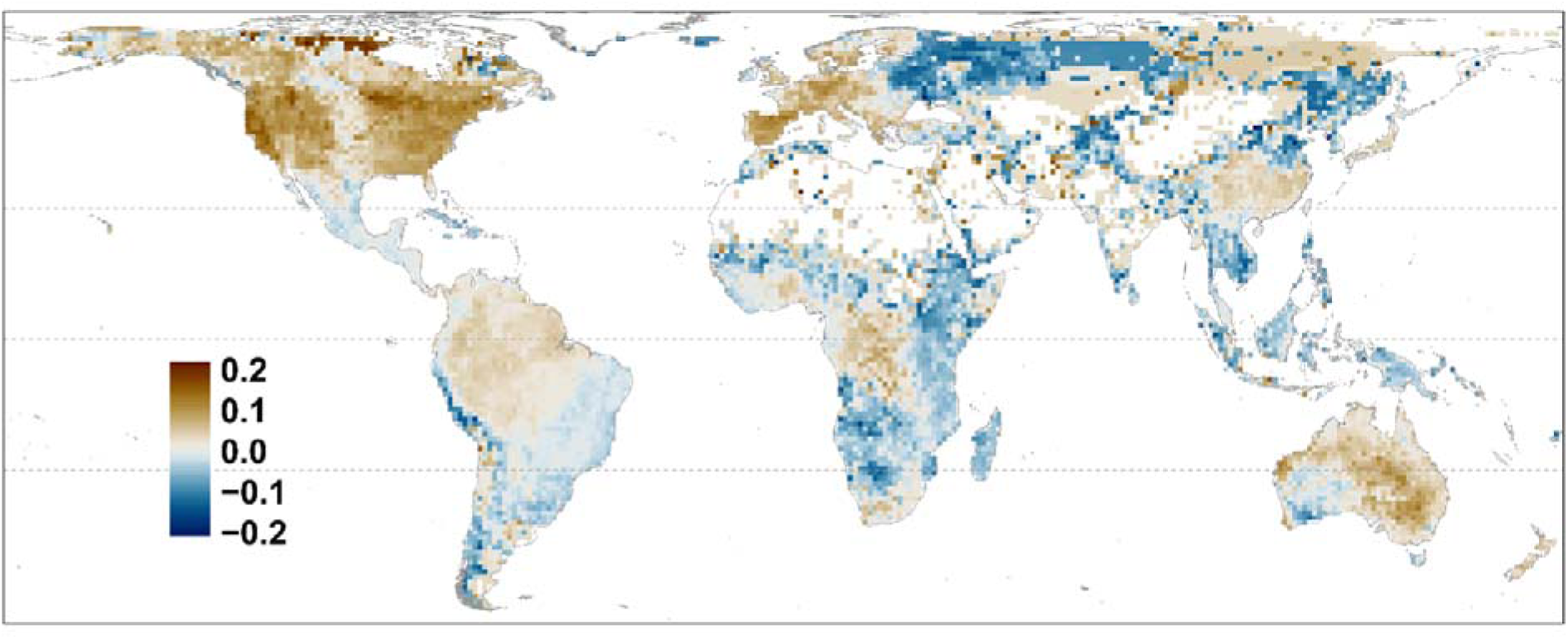
Global patterns of the residuals (deviation) from the ordinary least regression between functional diversity (FD) and phylogenetic diversity (PD) (FD = 0.90PD, *R*^2^ = 0.987, *p* < 0.0001). Brown (positive) areas are areas of higher FD than expected based on PD, whereas blue (negative) areas depict areas with lower FD than expected from the observed PD. Map uses the Behrmann projection at 110 km × 110 km spatial resolution.

The relative importance of the factors explaining variation in FD residuals are different from those explaining their variations (Fig. 4 vs. Fig. 2b & 2c). Overall, current AP is correlated negatively with the FD residuals both globally and regionally, but is only the strongest driver at global scale (Figs. 4 & 5, Table S3). MAT and non-climatic factors show weak or no relations, except for MAT for Indo-Maley and the Neotropics. The effects of the paleoclimate are variable. At global scale, the Miocene AP anomaly and the LGM MAT anomaly are negatively related to the FD residuals, while the LGM AP anomaly is positively related (Figs. 4 & 5). However, these relationships are inconsistent across biogeographic regions (Figs. 4 & 5).

**Fig. 4.**
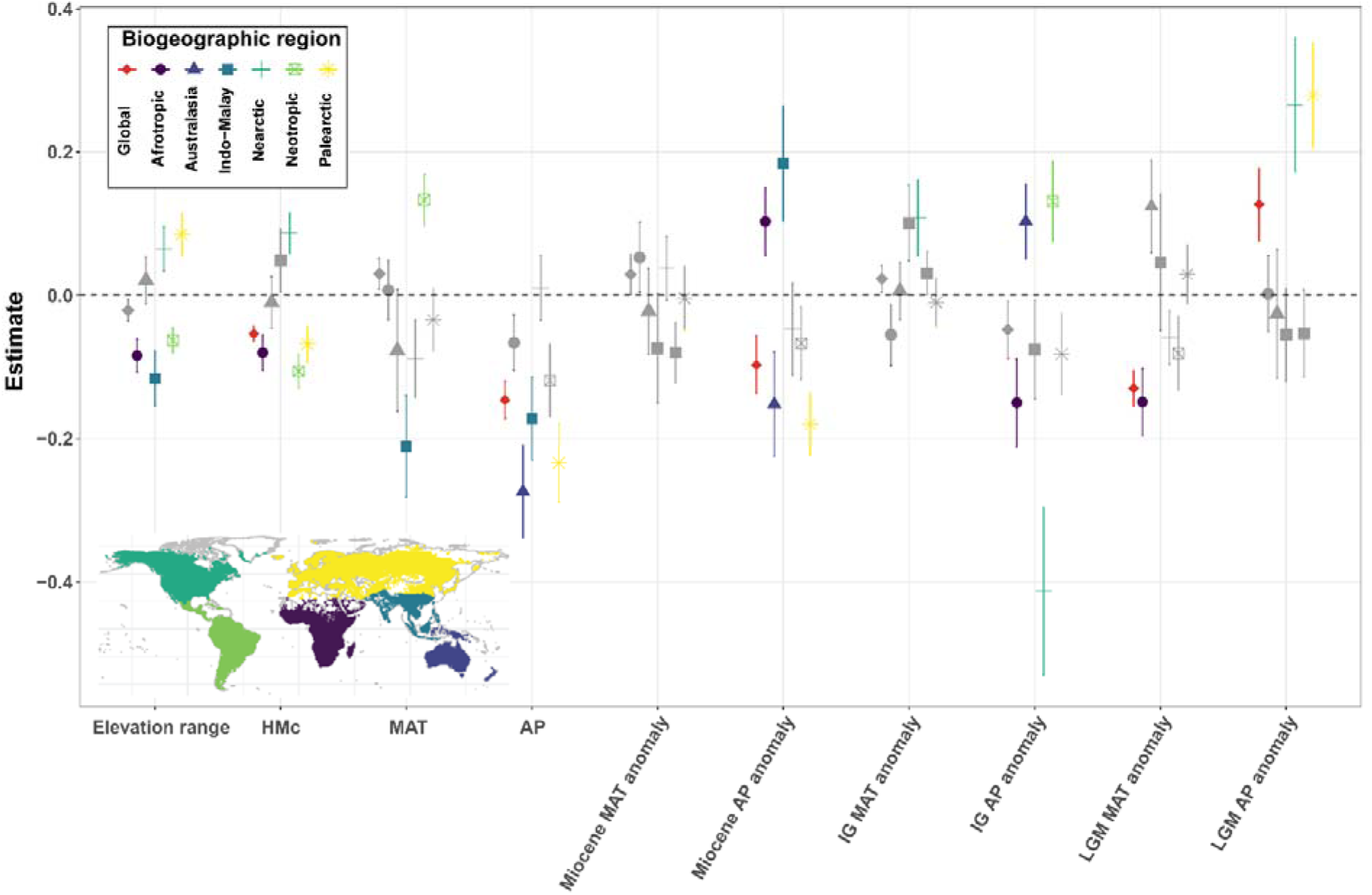
Effects of the tested environmental variables on the residuals from the regression between functional diversity (FD) and phylogenetic diversity (PD) (Fig. 3). Estimate (± 1standard error) of effects were obtained from simultaneous autoregressive (SAR) models. Different colors and shapes indicate biogeographic regions. Non-significant variables (*p* > 0.05) are indicated in grey. Results from OLS models are shown in Table S3. HMc: human modification index; MAT: mean annual temperature; AP: Annual precipitation; IG: Pleistocene Interglacial; LGM: Last Glacial Maximum. Anomaly was calculated as the past minus the present state.

**Fig. 5.**
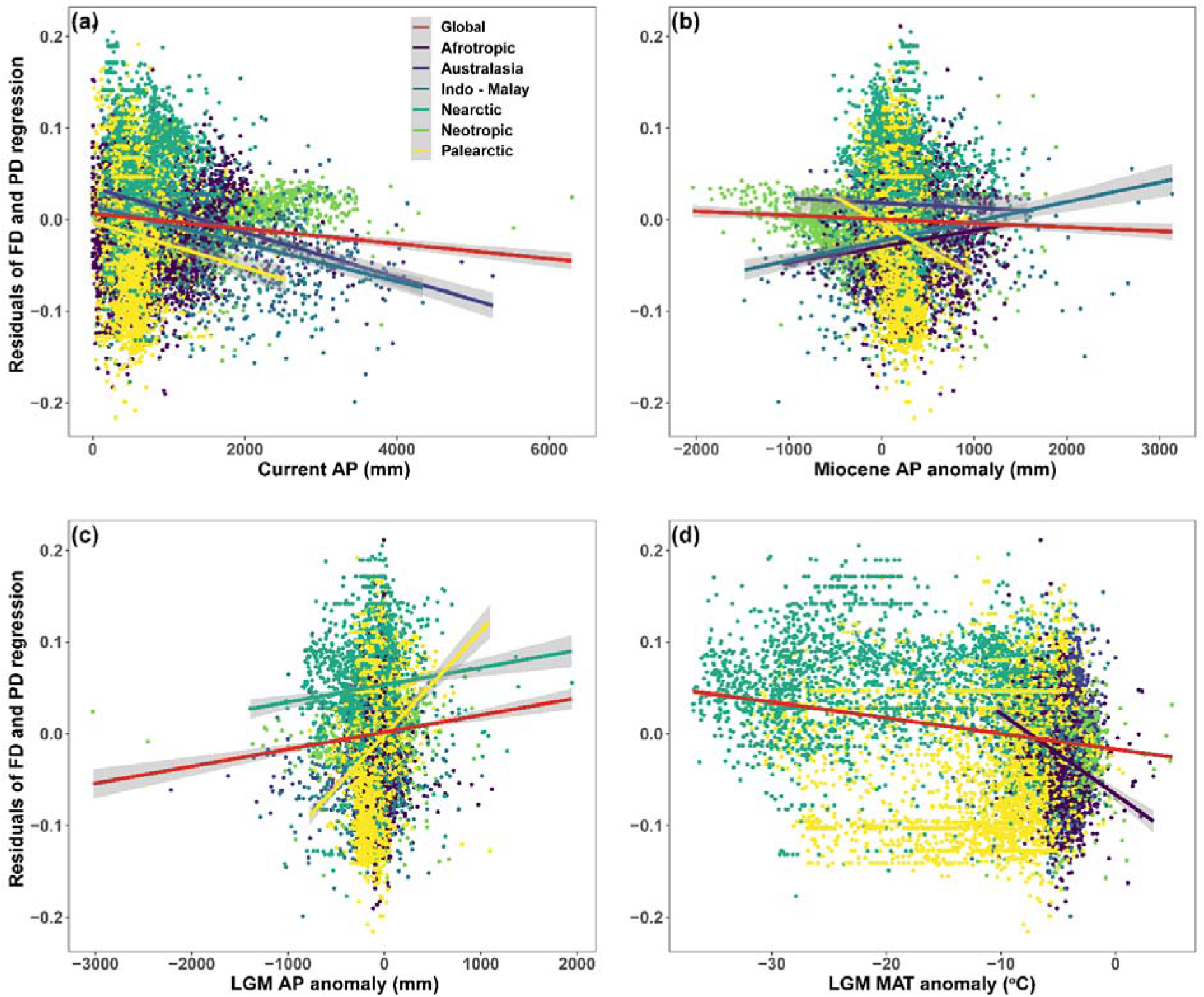
Bivariate regressions between the residuals from the regression of functional diversity (FD) on phylogenetic diversity (PD) and significant explanatory climate variables. Anomaly was calculated as the past minus the present state. For each subplot, only significant relationships are shown (Fig. 4, *p* < 0.05).

## Discussion

Based on an unprecedented tree occurrence database, our study maps strong latitudinal patterns in all three diversity dimensions (SR, PD, and FD) at global scale. The SR-linked global latitudinal patterns of Faith’s PD and FD matches previous empirical and modeled studies of tree species richness (e.g., ^9,61^), tree functional diversity in the New World ^62^, and tree phylogenetic diversity at a regional scale ^31^. It has been reported that speciation in rainforest environments has taken place at least since the Paleocene (∼58 mya) ^63,64^, probably coupled to jointly high temperatures and precipitation ^38,49,51,65^. Moreover, the relatively stable environment, compared to high latitudes, may also resulted in low extinction rates, making the tropics both “cradles (species diversifying)” and “museums (species persistence)” of species diversity ^49,66^. In addition, long speciation history and lower extinction rates in the tropics could result in both higher phylogenetic diversity and functional diversity ^38,48,49,51, but see 35^.

Our results provide evidence that paleoclimate complements current climate in shaping tree diversity globally and regionally, and that these effects are not only related to the recent prehistory – such as the Last Glacial period, represented by the LGM 21,000 years ago – but also much deeper time scales. These results extend previous findings for other organism groups notably for species richness and endemics ^39,42^ and for trees or plant clades including trees in specific regions and biomes ^26,27,29,67–70^, to trees globally. Importantly, they go beyond species richness to the more ecologically meaningful indices, functional and phylogenetic diversity.

Notably, we found that precipitation effects were stronger and more consistent (across regions) drivers than temperature effects, especially in relation to the wet and warm middle Miocene (11.6 – 7.5 mya), and the dry, cold LGM. The middle Miocene, the warmest and wettest interval in the late Cenozoic, was a period of forest expansion ^34,71^, due to warming coupled with elevated atmospheric CO_2_ (>500 ppm) ^72–74^. This likely promoted high species diversity globally due to a higher diversification rate and lower extinction rate ^38,48,51^. The Myrtaceae family ^75^ and the genus *Quercus* ^76^ are examples that follow this pattern. As a legacy of forest expansion, the generally warmer and wetter climate in the late Miocene compared to the contemporary climate have a positive associations to tree SR, FD, and PD ^37^. We also see this in our results at the global scale and for most regions with positive effects of both Miocene AP and MAT (Fig. 2). The weak negative association between the LGM MAT anomaly and SR, but not with FD and PD, could indicate that global cold climate in LGM (Fig. S1) caused range extractions or even extinctions of certain species. Likely, the intensity of these processes were not strong enough to significantly decrease the communities’ FD and PD, probably due to the high tree diversity accumulated in previous warm and humid periods ^36,75^. Indeed, both tree FD and PD showed the tendency to level-off with SR increase (Fig. S6), a similar pattern reported by ^54^, indicating that closely related tree species have more similar traits, i.e., the functional space tightly packed ^30,70^. The LGM precipitation anomaly was positively related to tree SR, PD and FD, likely reflecting widespread forest contractions during the generally dry LGM and tree survival in moist refugia ^77,78^. Furthermore, the diversity of drier forests itself is generally lower due to a limited number of niches and the physiological limits of species drought tolerance ^79^. Our results suggest that paleoclimate affects not just forest biodiversity, but also forest ecosystem functioning given the effects found here, which corroborates other studies on FD ^16,17^ and PD ^20,27,67^. Notably, a recent study has found that paleoclimatic legacies in tree FD negatively affect stand productivity in Northern Hemisphere temperate forests ^80^.

The relationships between paleoclimate and SR, FD, and PD were partially repeated within biogeographic regions, there was also substantial inter-region variation in these relations (Fig. 2). For example, not all of the four significant relationships found globally were retained regionally, and new relations emerged in some cases. These variable regional relations may reflect differing regional paleoclimatic histories, differences in the overall climatic and geographic setting, as well as methodological effects, e.g., different covariation among explanatory variables. For example, in Australasia, only the LGM AP anomaly showed significant, negative relationships with FD and PD, possibly because the temperature there was rather stably high during the last millions of years, with precipitation being more variable and lower (Fig. S2).

The regions representing FD surplus relative to PD, i.e., where species were found to be more functionally diverse (high FD) than expected from PD, largely coincided with high SR regions (Figs. 3 & 1a), represented by warm and humid climate today. This suggests that communities in warm and humid conditions have accumulated more FD than expected compared to dry or cold regions. This FD surplus could be caused by high competition, high heterogeneous environments, or otherwise diversifying trait evolution ^19,62,81–83^. We found that all precipitation variables were important for explaining the FD deviation from PD, even though their effects differed (Figs. 4 & 5). Surprisingly, high current precipitation tended to correspond to FD deficits, i.e., areas where species were more functionally similar than predicted by PD, both globally and in several biogeographic regions. Even though the observed FD in many wet and warm areas were higher than expected from PD, an explanation for the observed relationship could be that moist tropical forests harbor large numbers of shade-tolerant species, which have evolved along a similar evolutionary path (i.e., stabilizing selection) to adapt to the shady environment, thus showing high levels of ecological equivalence ^83,84^.

Building on recent progress in the harmonization of several databases on tree species distributions, functional traits, phylogenetic relatedness, and global paleoclimate, we have found that the tropics harbor the highest diversity across not only taxonomic, but also functional and phylogenetic dimensions, while high latitudes have lower diversity values for all diversity measures. Nevertheless, there are important and informative deviations between the patterns in FD and PD, including a signature consistent with less ecological filtering in moist, shady tropical forest environments ^84^. Importantly, we found evidence that current tree phylogenetic and functional diversities are likely shaped not only by the contemporary environment, but also by past climate as far back as the Miocene (∼10 Mya). Notably, we see long-term reductions in FD and PD in relation to past climatic cold or drought stress, likely affecting current forest ecosystem functioning ^80^. These findings highlight the importance of climate for tree diversity and forest ecosystems, and that losses from future climate change could have strong and very long-lasting effects.

## Methods

### Tree species and their range maps

In this study, we used the world tree species list ^56^ and species range maps compiled by ^47,57^. Briefly, the world tree species checklist (GlobalTreeSearch, GTS ^56^) was used to extract the global tree species list for the current study. Tree species included in the GTS is based on the definition by the IUCN’s Global Tree Specialist Group (GTSG), i.e., “a woody plant with usually a single stem growing to a height of at least two meters, or if multi-stemmed, then at least one vertical stem five centimeters in diameter at breast height” ^56^. This list was subsequently standardized via the Taxonomic Name Resolution Service (TNRS) online tool ^85^ to remove synonyms. The occurrence records of the selected species were collated from five widely used and publicly accessible databases, namely: the Global Biodiversity Information Facility (GBIF; http://www.gbif.org), the public domain Botanical Information and Ecological Network v.3 (BIEN; http://bien.nceas.ucsb.edu/bien/;^86,87^), the Latin American Seasonally Dry Tropical Forest Floristic Network (DRYFLOR; http://www.dryflor.info/; ^88^), the RAINBIO database (http://rainbio.cesab.org/; ^89^), and the Atlas of Living Australia (ALA; http://www.ala.org.au/). The compiled occurrence data was accessed ^57^ and the high-quality records were then used to generate range maps based on the alpha hull algorithm via the *Alphahull* package ^90,91^ in R (ver. 3.5.1; ^92^). We further validated the range maps using an external independent dataset ^9^. The estimated range maps of the 46,752 tree species were rasterized to 110 km equal-area grid cells (∼1 degree at the Equator), a resolution commonly used in global diversity studies (e.g.,^45^), using the *letsR* package ^93^. For detailed information on the range map estimations and external validation, see ^47^.

### Phylogeny

We constructed a phylogenetic tree for the tree species using the largest seed-plant phylogeny presently available (the ALLMB tree ^94^). This dated phylogeny combines a backbone tree ^95^, which was built using sequence data from public repositories (GenBank) to reflect deep relationships, with previous knowledge of phylogenetic relationships and species names from the Open Tree of Life (Open Tree of Life synthetic tree release 9.1 and taxonomy version 3, https://tree.opentreeoflife.org/about/synthesis-release/v9.1). This phylogeny was matched to our tree species dataset, and any species that were not in our dataset were removed from the tree. Subsequently, some species missing from the phylogeny were manually added, using the same approach as ref. ^94^.

### Functional trait data

Eight ecologically relevant and commonly used traits ^96^ were selected for functional diversity analyses, i.e., leaf nitrogen content, wood density, leaf phosphorus content, leaf dry matter content, plant max height, seed dry mass, specific leaf area, and leaf area. Originally, we compiled 21 functional traits from the TRY (https://try-db.org/TryWeb/Home.php; ^97,98^, TOPIC ^99–105^, and BIEN (http://bien.nceas.ucsb.edu/bien/; ^86,87^) databases. As many of the species’ trait were missing, we imputed missing values via an gap-filling algorithm with Bayesian Hierarchical Probabilistic Matrix Factorization (BHPMF, ^106–108^), which is mostly based on both trait-trait correlation matrix and the phylogentic signal of traits (Refer to ref. ^47^ for the detailed gap-filling procedure). In this process, all the 21 traits were used to maximally benefit from the correlations among them.

### Environmental variables

We compiled 17 environmental variables, including current climate, paleo-climate, human effects, topographic heterogeneity and evolutionary history (Supplementary Table S1). Climate, both present-day and paleoclimate, is generally assumed to be a vital predictor of species distribution and diversity patterns (e.g., ^26,27,29,39,109,110^). Due to the data availability of the paleoclimates, we included two bioclimatic predictors commonly used in relevant studies: annual mean temperature (MAT) and annual precipitation (AP). Current climate variables were extracted from WorldClim (v.2, www.worldclim.org) at a resolution of 30 arc-seconds (∼1 km at the equator), averaging global climate data from the period 1970 - 2000 ^111^. We selected six paleo-time periods spanning from *ca*. 11.6 – 7.2 mya to *ca*. 21 kya, representing climatic conditions either warmer, cooler, or similar compared to the present-day climate. Specifically, each bioclimatic layer of the late Miocene climate (11.61 – 7.25 mya ^37^) and mid-Pliocene Warm period (∼ 3.264 – 3.025 mya; ^112,113^) were averaged to represent the warmer climate compared to present day (hereafter Miocene). Pliocene Marine Isotope Stage M2, a glacial interval in the late Pliocene (∼ 3.3□mya; ^113,114^), was used to represent the Pliocene global cooling period, while the Last Glacial Maximum (LGM, ∼ 21 kya) was used to present the more recent global cooling event compared to M2 ^113,115^. We further constructed a current climate (hereafter Interglacial, IG) analog using the mean value per bioclimatic layer between the Pleistocene Marine Isotope Stage 19 (MIS 19), the oldest Pleistocene interglacial (∼ 787 kya ^113^), and the Last Interglacial (LIG; ∼ 130 kya ^116^). The mid-Pliocene Warm Period, Pliocene M2, Pleistocene MIS19, and the LIG data were extracted from Paleoclim (www.paleoclim.org), at a resolution of 2.5 arc-minutes (∼ 4.5 km at the equator) ^113^, and the LGM data was extracted from the CHELSA database (www.chelsa-climate.org) at a resolution of 30 secs ^115^.

In addition to climate, other factors, such as human activities, topographic heterogeneity, and evolutionary history, can also affect plant distributions ^9,26,117,118^. The Human Modification map (HMc ^119^)^119^ was used as a proxy of human activities. Compared to the commonly used human footprint index and human influence index maps ^120^, HMc has been modelled with the incorporation of 13 most recent global-scale anthropogenic layers (with the median year of 2016) to account for the spatial extent, intensity, and co-occurrence of human activities, many of which showing high direct or indirect impact on biodiversity ^121^. HMc was extracted at a resolution of 1 km^2 119^. The elevation range is the absolute difference between the maximum and minimum elevation value within a specific area. We computed the elevation/topographic range within each 110 ×110 km grid cell based on the digital elevation model at 90 m resolution (http://srtm.csi.cgiar.org/). Elevation range is a proxy of environmental heterogeneity, which is considered as a universal driver of biological diversity ^122,123^. To analyze the potential effects of evolutionary and biogeographic history, we also included the biogeographic regions as an additional variable. We applied the definition of biogeographic regions from ref. ^124^, which defines 12 regions globally using cladistic and phylogenetic analyses of plant species, and plate tectonics. However, due to the varying data size in each of the 12 regions, we combined them into six regions, i.e., Afrotropic, Australasia, Indo-Malay, Nearctic, Neotropic, Palearctic, largely similar to the biogeographic realms proposed by ref. ^125^. All predictors were extracted from various databases, which we describe in further detail in the supplement (Supplementary Table S1).

Except for the biogeographic regions and elevation range, mean values for all predictors were extracted at a 110 ×110 km resolution. The variable extractions and averaging were carried out in the *letsR* package. Due to the low reliability and/or missing environmental variables for many islands ^126^, we removed insular grid cells from small islands, and 11,950 grid cells with records were kept (Fig. S5).

### Phylogenetic and functional diversity

Phylogenetic diversity (PD) was calculated for each 110 ×110 km grid cell as the sum of the branch lengths of all co-occurring species as defined by ref. ^10^. Among the many existing, somewhat overlapping matrices of PD, the one we selected is the most widely used due to its easy calculation and interpretation and a more robust basis for conservation ^10,13,14^.

Functional diversity (FD) was calculated in an analogous manner to PD ^127^. A Principal Component Analysis (PCA) was applied to the eight traits to eliminate trait redundancy. Values of all traits were log transformed to improve normality and were standardized before analysis. Then a dendrogram based on the first three PCs (explaining 84% of the total variation) was constructed using Gower’s distance via the *vegan* ^128^ and *fastcluster* ^129^ packages. This dendrogram was used to calculate FD as the sum of the total branch lengths connecting a set of species in the 110 ×110 km grid cell. Both PD and FD were calculated using the *letsR* and *picante* ^130^ packages.

To investigate the bivariate relationships between FD and PD, an ordinary least squares model was implemented. We further plotted the residuals of model to show any deviation between FD and PD.

### Statistical analyses

To test the long-term climate stability hypothesis, we calculated the anomaly for MAT and AP between the four paleo-time periods and the present-day, i.e., past minus present, to represent the amplitude of the climate changes within each time-scale (Fig. S1) ^26,27,29,39,118^. On average, compared to the present, mean annual temperature (MAT) was much higher in the Miocene, slightly higher in the Pliocene M2 period, much lower in the LGM, and similar in the IG (Fig. S2a). During Pliocene M2 and IG, annual precipitation (AP) was similar to the present-day, while the Miocene and LGM had slightly higher or lower precipitation, respectively than the contemporary precipitation (Fig. S2b). The paleo-time periods selected, thus, represent (on average) cold, warm, and similar paleo-climates compared to present-day conditions.

Pearson correlation coefficients showed a low level of correlations between MAT, AP, and their respective anomaly variables (Fig. S6). However, MAT and AP of Pliocene M2 and Pleistocene IG anomaly showed relatively high correlations (Fig. S7) with or without accounting for the spatial autocorrelation (using the *SpatialPack* package ^131^). Consequently, we removed the two Pliocene M2 variables from further analyses.

We used ordinary least squares models (OLSs) and simultaneous autoregressive models (SARs), if the OLS model residuals exhibited spatial autocorrelation (SAC), to evaluate the relative importance of the predictor variables in determining the variation in each of the three diversity indices and the residuals of bivariate relationships between FD and PD. We used the SAR error model because of its superior performance compared to other SAR model types ^132^. The SAR error model adds a spatial weights matrix to an OLS regression to accounts for SAC in the model residuals. A series of spatial weights, i.e., *k*-means neighbor of each site, were tested and *k* = 1.5 was used for all SARs models as it can successfully account for the SAC (see Supplementary results of statistical analyses). Residual SAC was examined in all models (both OLS and SAR) using Moran’s *I* test, and Moran’s *I* correlograms were also used to visualize the spatial residuals of the models. Model explanatory power was represented by adjusted *R*^2^ (OLSs) and Nagelkerke pseudo-*R*^2^ (SARs) ^133^, while the Akaike Information Criterion (AIC) and Bayesian information criterion (BIC) were used to compare the models for each diversity index ^134^. SARs and Moran’s *I* tests were carried out using the *spdep* package ^135^. Both OLS and SAR models were run by including current MAT and AP, the six anomaly variables, and the other non-climate predictors (elevation range and HMc) to investigate their relative contributions to each diversity index. In addition to the global models, we ran the same models for each biogeographic region to test whether the global relationships varied among regions. Moreover, we ran three additional global models for the FD and PD indices, selecting only one paleoclimate (both MAT and AP) from the three paleo-time periods at the time, and keeping other variables the same in each model to investigate whether the effects of the different paleoclimate predictors changed compared to the full models (including all paleo climatic predictors). Before running the models, we inspected the normality of all predictors and log_10_-transformed variables if needed. All response variables (three diversity indices) were log_10_-transformed. Thereafter, we standardized all predictor variables by transforming all variables to a mean of zero and a standard deviation of one to derive more comparable estimates ^136^.

### Supplementary results of statistical analyses

We found that for all models (both global and regional), SAR models performed better than the corresponding OLS models, regarding to AIC, BIC, and *R*^2^ (Tables S2-S3), and all SAR models successfully accounted for SAC in model residuals (*p* >> 0.05, Figs. S8-S11). Thus, we only represented the results from SARs models in the text, even though the significance of some predictors varied between OLS and SAR models (Fig. S12). In addition, we found that the effects of paleoclimate variables showed no change between the full models, including all paleoclimate variables and models using paleoclimate of each paleo-period (Fig. S13-S14). This clearly shows the robustness of their relationships with the tree diversity indices.

## Acknowledgments

We thank Brad Boyle for valuable database and informatics assistance and advice, and TRY contributors for sharing their data. This work was conducted as a part of the BIEN Working Group, 2008–2012. We thank all the data contributors and numerous herbaria who have contributed their data to various data compiling organizations (see the herbarium list below) for the invaluable data and support provided to BIEN. We thank the New York Botanical Garden; Missouri Botanical Garden; Utrecht Herbarium; the UNC Herbarium; and GBIF, REMIB, and SpeciesLink. The staff at CyVerse provided critical computational assistance.

We acknowledge the herbaria that contributed data to this work: A, AAH, AAS, AAU, ABH, ACAD, ACOR, AD, AFS, AK, AKPM, ALCB, ALTA, ALU, AMD, AMES, AMNH, AMO, ANGU, ANSM, ANSP, AQP, ARAN, ARIZ, AS, ASDM, ASU, AUT, AV, AWH, B, BA, BAA, BAB, BABY, BACP, BAF, BAFC, BAI, BAJ, BAL, BARC, BAS, BBB, BBS, BC, BCMEX, BCN, BCRU, BEREA, BESA, BG, BH, BHCB, BIO, BISH, BLA, BM, BOCH, BOL, BOLV, BONN, BOON, BOTU, BOUM, BPI, BR, BREM, BRI, BRIT, BRLU, BRM, BSB, BUT, C, CALI, CAN, CANB, CANU, CAS, CATA, CATIE, CAY, CBM, CDA, CDBI, CEN, CEPEC, CESJ, CGE, CGMS, CHAM, CHAPA, CHAS, CHR, CHSC, CIB, CICY, CIIDIR, CIMI, CINC, CLEMS, CLF, CMM, CMMEX, CNPO, CNS, COA, COAH, COCA, CODAGEM, COFC, COL, COLO, CONC, CORD, CP, CPAP, CPUN, CR, CRAI, CRP, CS, CSU, CSUSB, CTES, CTESN, CU, CUVC, CUZ, CVRD, DAO, DAV, DBG, DBN, DES, DLF, DNA, DPU, DR, DS, DSM, DUKE, DUSS, E, EA, EAC, EAN, EBUM, ECON, EIF, EIU, EMMA, ENCB, ER, ERA, ESA, ETH, F, FAA, FAU, FAUC, FB, FCME, FCO, FCQ, FEN, FHO, FI, FLAS, FLOR, FM, FR, FRU, FSU, FTG, FUEL, FULD, FURB, G, GAT, GB, GDA, GENT, GES, GH, GI, GLM, GMDRC, GMNHJ, GOET, GRA, GUA, GZU, H, HA, HAC, HAL, HAM, HAMAB, HAO, HAS, HASU, HB, HBG, HBR, HCIB, HEID, HGM, HIB, HIP, HNT, HO, HPL, HRCB, HRP, HSC, HSS, HU, HUA, HUAA, HUAL, HUAZ, HUCP, HUEFS, HUEM, HUFU, HUJ, HUSA, HUT, HXBH, HYO, IAA, IAC, IAN, IB, IBGE, IBK, IBSC, IBUG, ICEL, ICESI, ICN, IEA, IEB, ILL, ILLS, IMSSM, INB, INEGI, INIF, INM, INPA, IPA, IPRN, IRVC, ISC, ISKW, ISL, ISTC, ISU, IZAC, IZTA, JACA, JBAG, JBGP, JCT, JE, JEPS, JOTR, JROH, JUA, JYV, K, KIEL, KMN, KMNH, KOELN, KOR, KPM, KSC, KSTC, KSU, KTU, KU, KUN, KYO, L, LA, LAGU, LBG, LD, LE, LEB, LIL, LINC, LINN, LISE, LISI, LISU, LL, LMS, LOJA, LOMA, LP, LPAG, LPB, LPD, LPS, LSU, LSUM, LTB, LTR, LW, LYJB, LZ, M, MA, MACF, MAF, MAK, MARS, MARY, MASS, MB, MBK, MBM, MBML, MCNS, MEL, MELU, MEN, MERL, MEXU, MFA, MFU, MG, MGC, MICH, MIL, MIN, MISSA, MJG, MMMN, MNHM, MNHN, MO, MOL, MOR, MPN, MPU, MPUC, MSB, MSC, MSUN, MT, MTMG, MU, MUB, MUR, MVFA, MVFQ, MVJB, MVM, MW, MY, N, NA, NAC, NAS, NCU, NE, NH, NHM, NHMC, NHT, NLH, NM, NMB, NMNL, NMR, NMSU, NSPM, NSW, NT, NU, NUM, NY, NZFRI, O, OBI, ODU, OS, OSA, OSC, OSH, OULU, OWU, OXF, P, PACA, PAMP, PAR, PASA, PDD, PE, PEL, PERTH, PEUFR, PFC, PGM, PH, PKDC, PLAT, PMA, POM, PORT, PR, PRC, PRE, PSU, PY, QCA, QCNE, QFA, QM, QRS, QUE, R, RAS, RB, RBR, REG, RELC, RFA, RIOC, RM, RNG, RSA, RYU, S, SACT, SALA, SAM, SAN, SANT, SAPS, SASK, SAV, SBBG, SBT, SCFS, SD, SDSU, SEL, SEV, SF, SFV, SGO, SI, SIU, SJRP, SJSU, SLPM, SMDB, SMF, SNM, SOM, SP, SPF, SPSF, SQF, SRFA, STL, STU, SUU, SVG, TAES, TAI, TAIF, TALL, TAM, TAMU, TAN, TASH, TEF, TENN, TEPB, TEX, TFC, TI, TKPM, TNS, TO, TOYA, TRA, TRH, TROM, TRT, TRTE, TU, TUB, U, UADY, UAM, UAMIZ, UB, UBC, UC, UCMM, UCR, UCS, UCSB, UCSC, UEC, UESC, UFG, UFMA, UFMT, UFP, UFRJ, UFRN, UFS, UGDA, UH, UI, UJAT, ULM, ULS, UME, UMO, UNA, UNB, UNCC, UNEX, UNITEC, UNL, UNM, UNR, UNSL, UPCB, UPEI, UPNA, UPS, US, USAS, USF, USJ, USM, USNC, USP, USZ, UT, UTC, UTEP, UU, UVIC, UWO, V, VAL, VALD, VDB, VEN, VIT, VMSL, VT, W, WAG, WAT, WELT, WFU, WII, WIN, WIS, WMNH, WOLL, WS, WTU, WU, XAL, YAMA, Z, ZMT, ZSS, and ZT.

W.-Y.G., J.M.S.-D., and J.-C.S. acknowledge support from the Danish Council for Independent Research | Natural Sciences (Grant 6108-00078B) to the TREECHANGE project. J.-C.S. also considers this work a contribution to his VILLUM Investigator project “Biodiversity Dynamics in a Changing World” funded by VILLUM FONDEN. C.B. was supported by the National Research Foundation of Korea (NRF) grant funded by the Korean government (MSIT) (2018R1C1B6005351). A.S.M. was supported by the Environment Research and Technology Development Fund (S-14) of the Ministry of the Environment, Japan. J.P. (Jan Pisek) was supported by the Estonian Research Council grant PUT1355. J.P. (Josep Peñuelas) was funded by the European Research Council Synergy grant ERC-2013-SyG-610028 IMBALANCE-P. A.G.G. (Alvaro G. Gutiérrez) was funded by FONDECYT 11150835 and 1200468. The BIEN working group was supported by the National Center for Ecological Analysis and Synthesis, a center funded by NSF EF-0553768 at the University of California, Santa Barbara and the State of California. Additional support for the BIEN working group was provided by iPlant/CyVerse via NSF DBI-0735191. B.J.E. and C.M. were supported by NSF ABI-1565118 and NSF HDR-1934790. B.J.E. was also supported by the Global Environment Facility SPARC project grant (GEF-5810). B.J.E., C.V., and B.S.M. acknowledge the FREE group funded by the synthesis center CESAB of the French Foundation for Research on Biodiversity (FRB) and EDF. J.-C.S. and B.J.E. acknowledge support from the Center for Informatics Research on Complexity in Ecology (CIRCE), funded by the Aarhus University Research Foundation under the AU Ideas program.

## Author contributions

W.-Y.G., J.M.S.-D., and J.-C.S. conceived the project; J.M.S.-D., W.-Y.G., and all others collected the data; W.-Y.G. analyzed the data; W.-Y.G. interpreted the data; W.-Y.G., J.M.S.-D., and J.-C.S. wrote the manuscript. All authors contributed data, discussed the results, revised manuscript drafts, and contributed to writing and approved the final manuscript.

## Data and materials availability

All the occurrences are deposited in BIEN (https://bien.nceas.ucsb.edu/bien/), and the phylogeny and imputed functional trait data are available via ref. ^47^.

