## Supplementary material for "Paleoclimate and current climate collectively shape the phylogenetic and functional diversity of trees worldwide": Supplemantary material

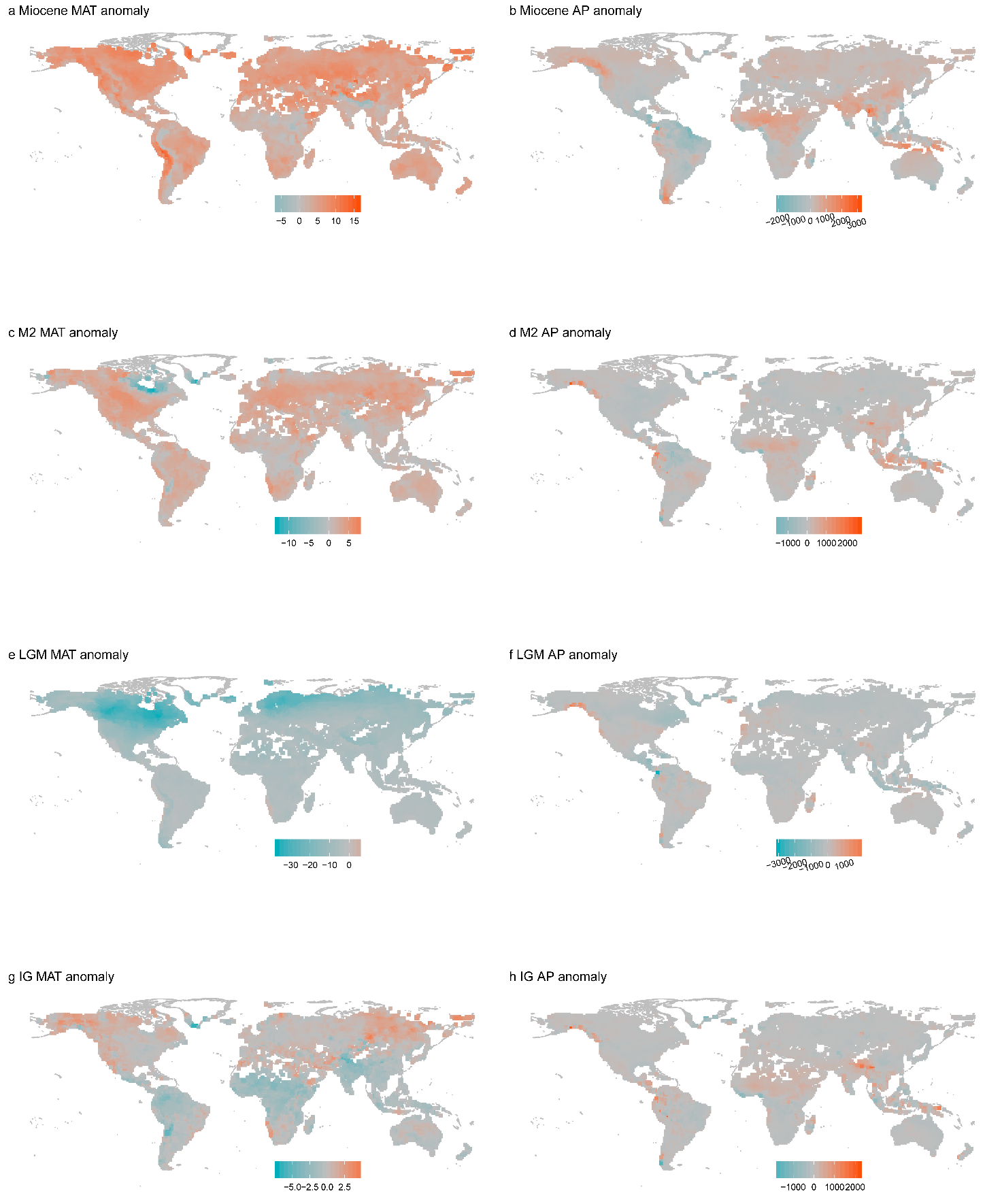
**Fig. S1** Maps of the MAT and AP anomaly in each paleo-period.

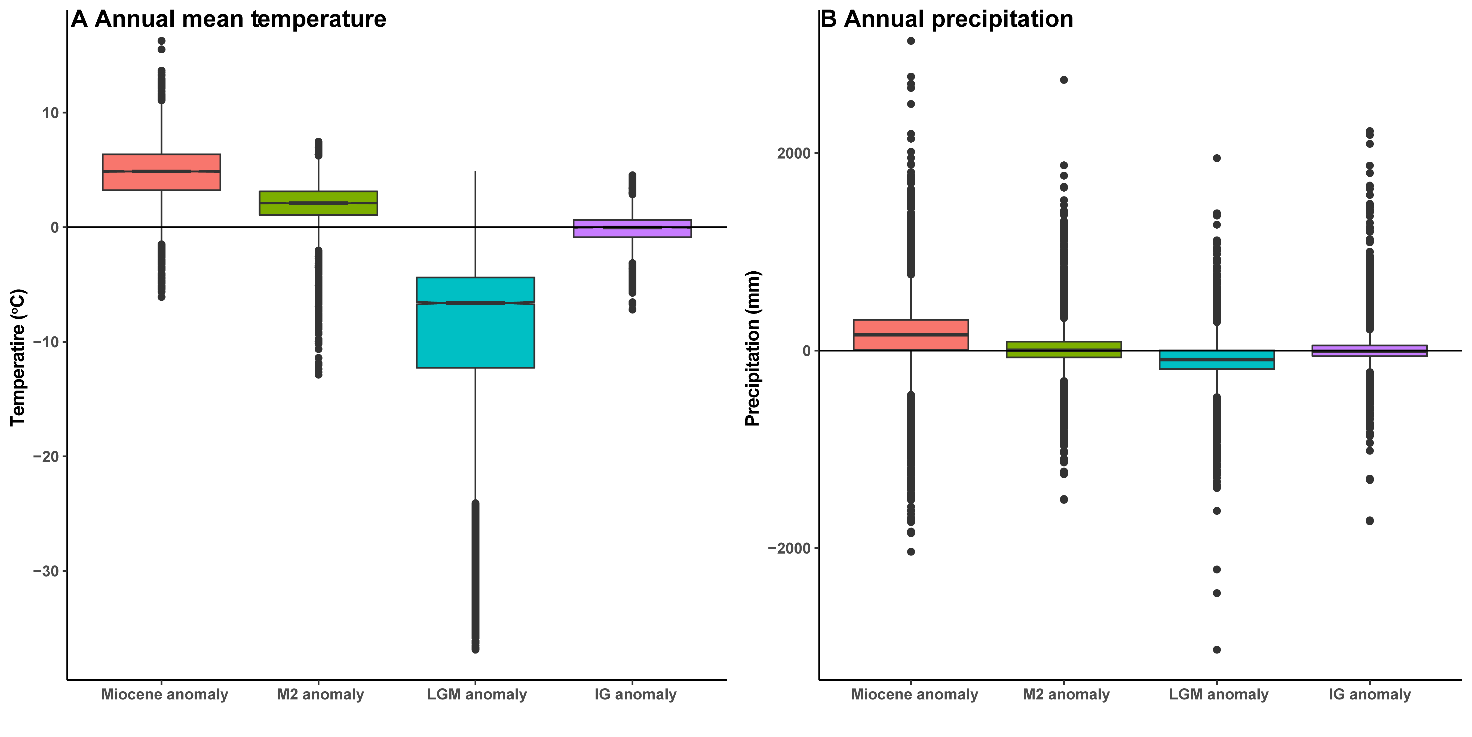

**Fig. S2** Boxplots of the temperature anomaly (A) and precipitation anomaly (B). Anomaly was calculated as the difference between the paleo-climate variable and the corresponding current climate variable (e.g., LGM MAT – current MAT). M2: Pliocene Marine Isotope Stage (MIS) M2. LGM: Last Glacial Maximum; IG: Pleistocene Interglacial.

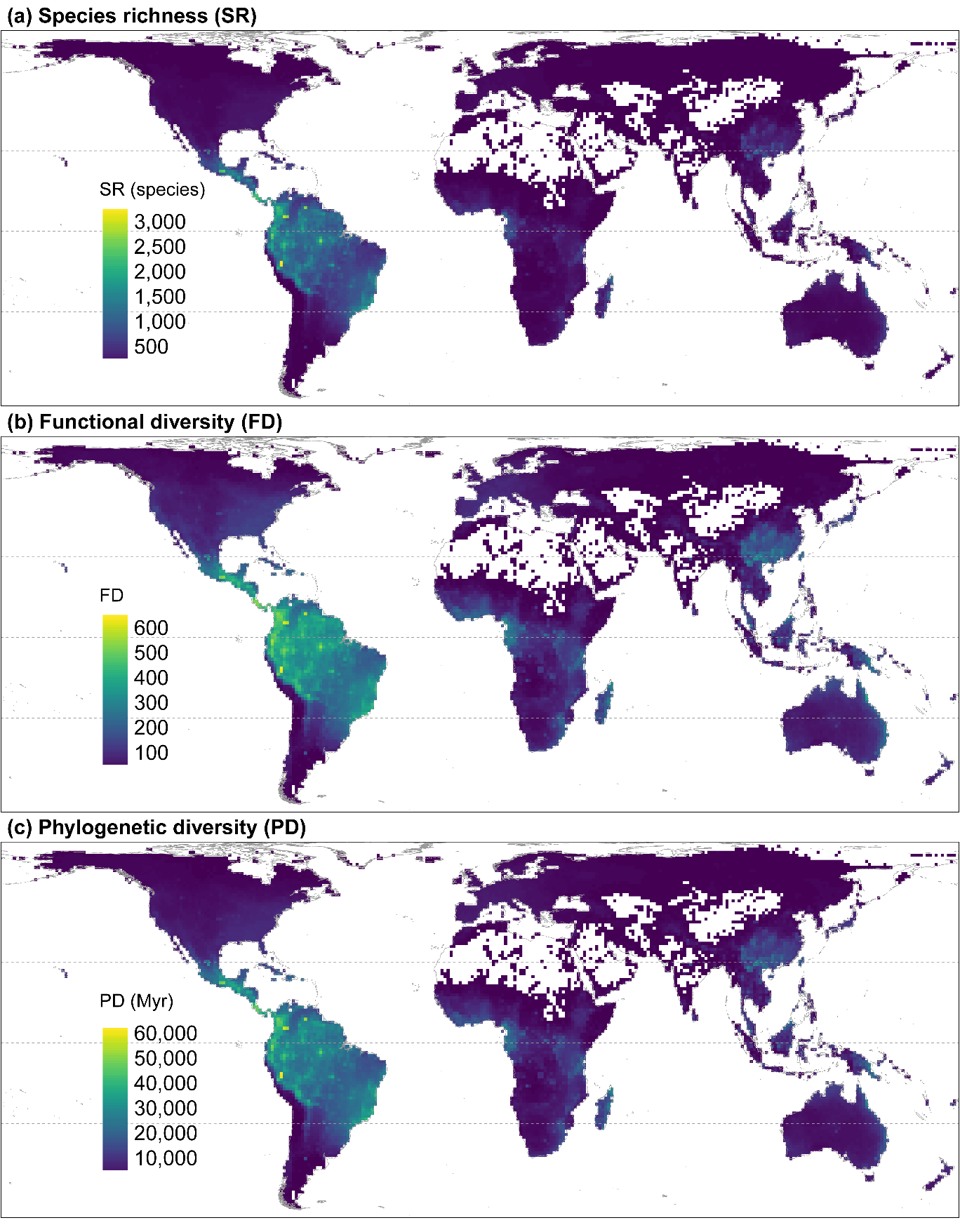

**Fig. S3** Global patterns of tree (a) species richness, (c) functional diversity, and (e) phylogenetic diversity. Maps use the Behrmann projection at 110 km × 110 km spatial resolution. Myr: Million years.

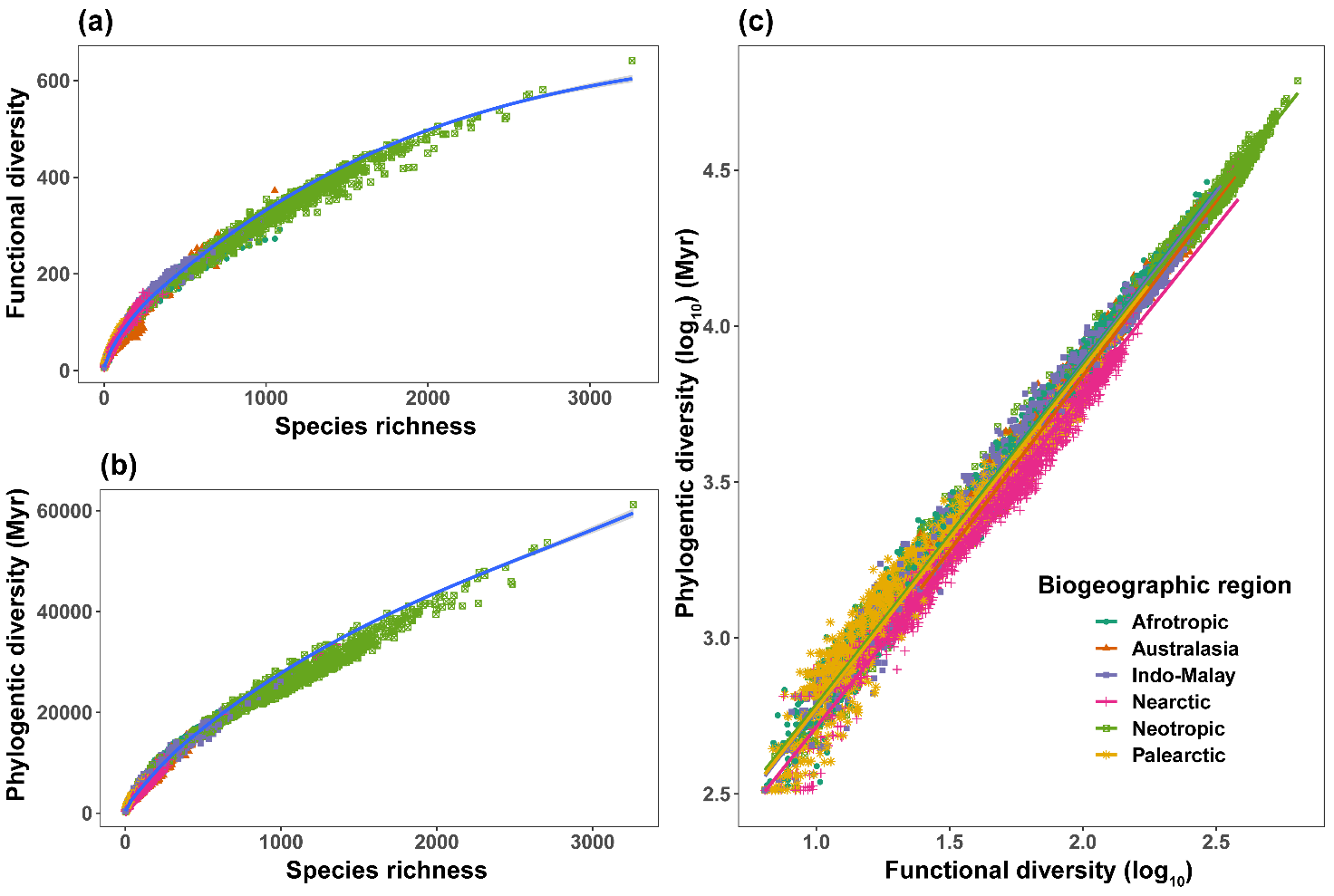

**Fig. S4** Relationships between (a) species richness and functional diversity, (b) species richness and phylogentic diversity, and (c) functional diversity and phylogenetic diversity for different biogeographic regions. For all the three relationships, *p* < 0.0001.

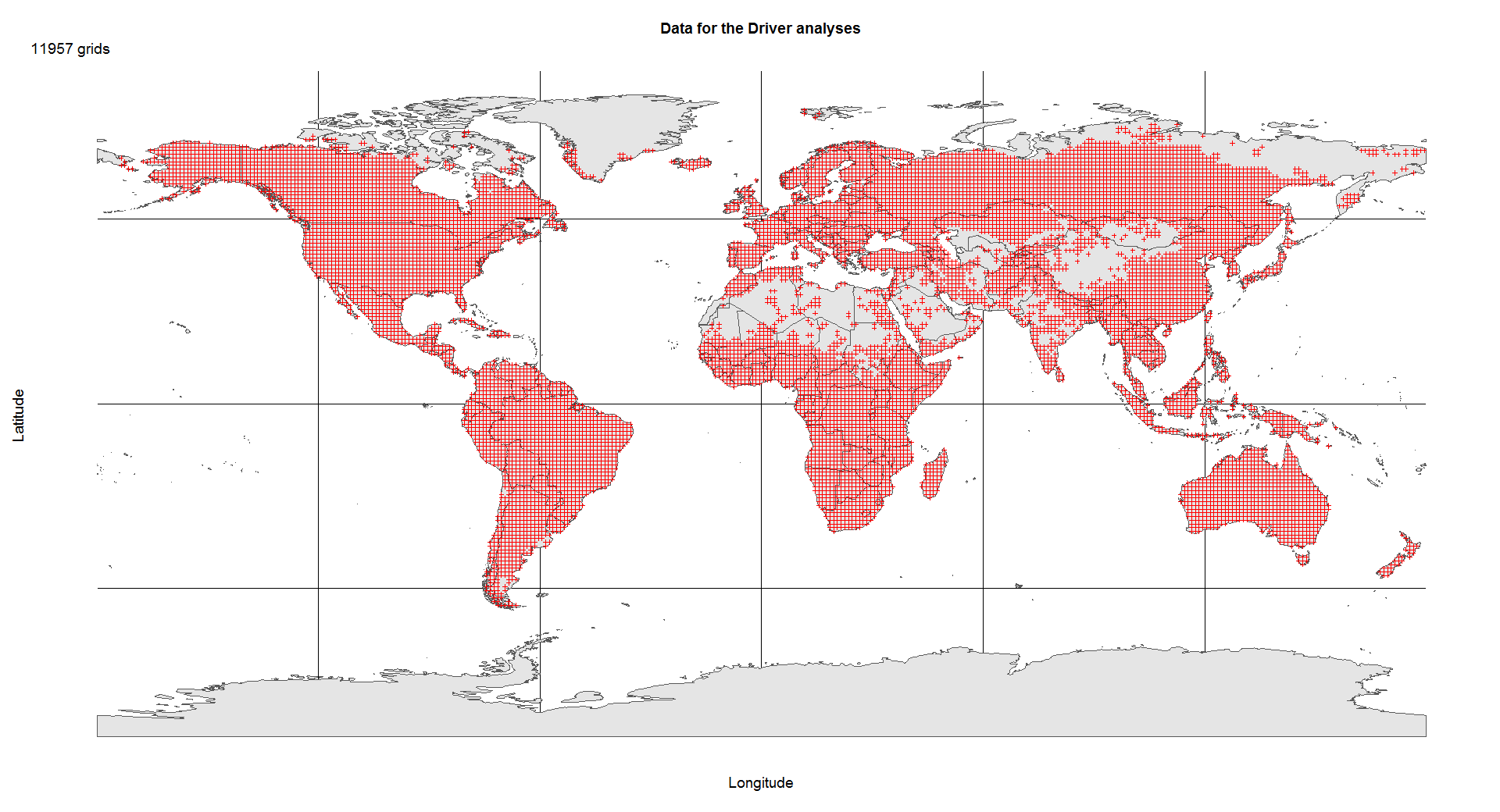

**Fig. S5** Data used in the driver analysis section. Overall, there were 11,950 grid cells with sufficient environmental data.

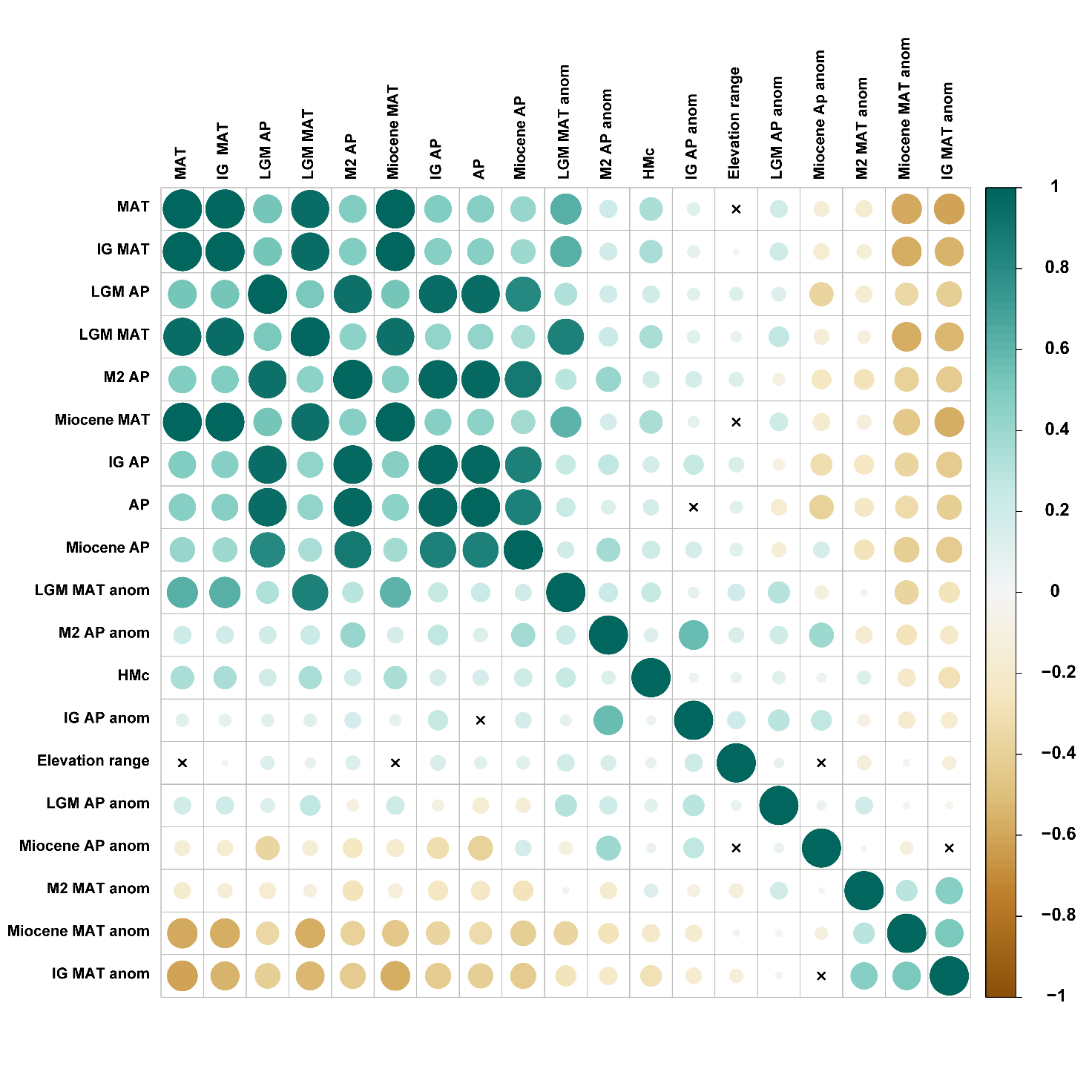

**Fig. S6** Correlation matrix among anomaly variables and other potential drivers. MAT: mean annual temperature. AP: Annual precipitation; HMc: human modification index. IG: Pleistocene Interglacial; LGM: Last Glacial Maximum; M2: Pliocene Marine Isotope Stage (MIS) M2; anom: anomaly.

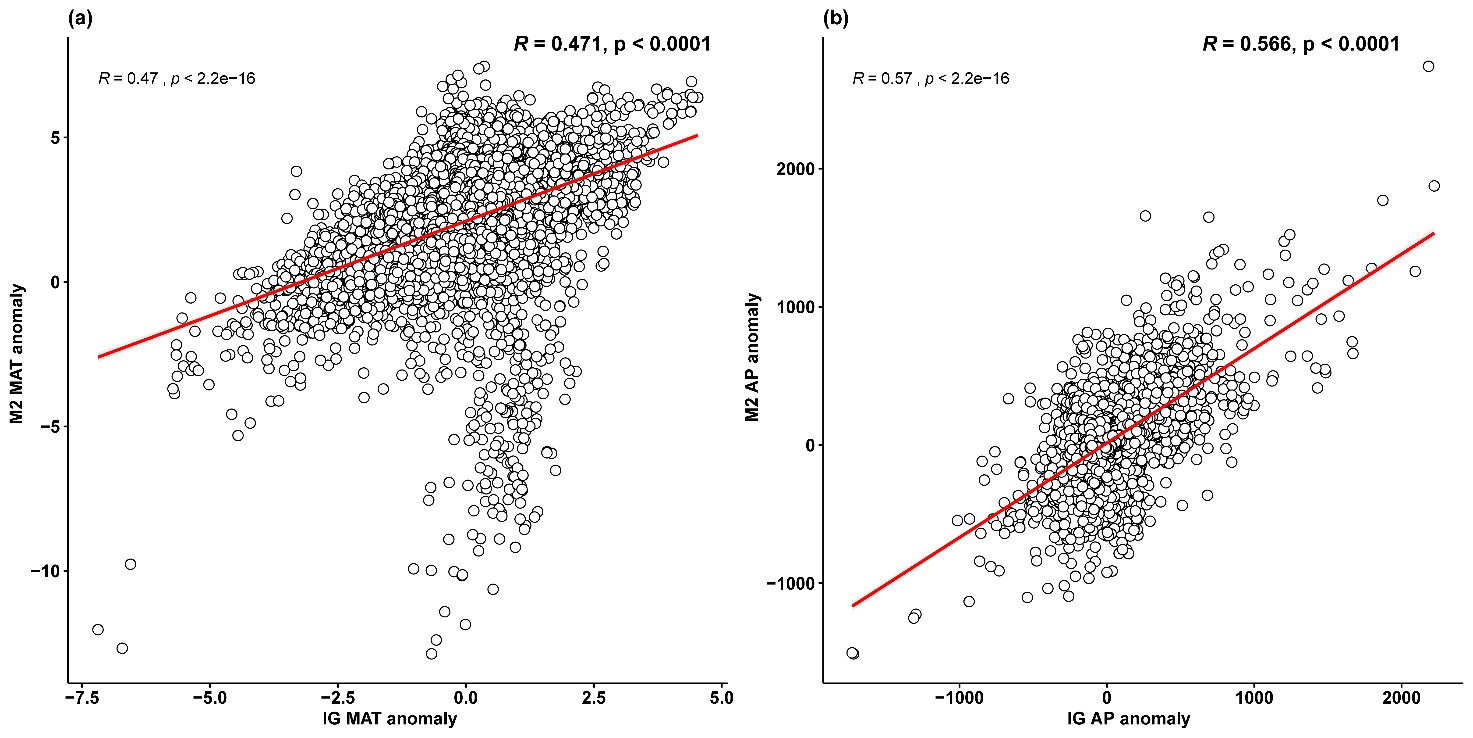

**Fig. S7** Relationships between Pliocene M2 and Pleistocene interglacial anomaly variables. The spatial adjusted *p*-values for the regression were shown at the top right of each subfigure. IG: Pleistocene Interglacial; M2: Pliocene Marine Isotope Stage (MIS) M2; MAT: mean annual temperature. AP: Annual precipitation.

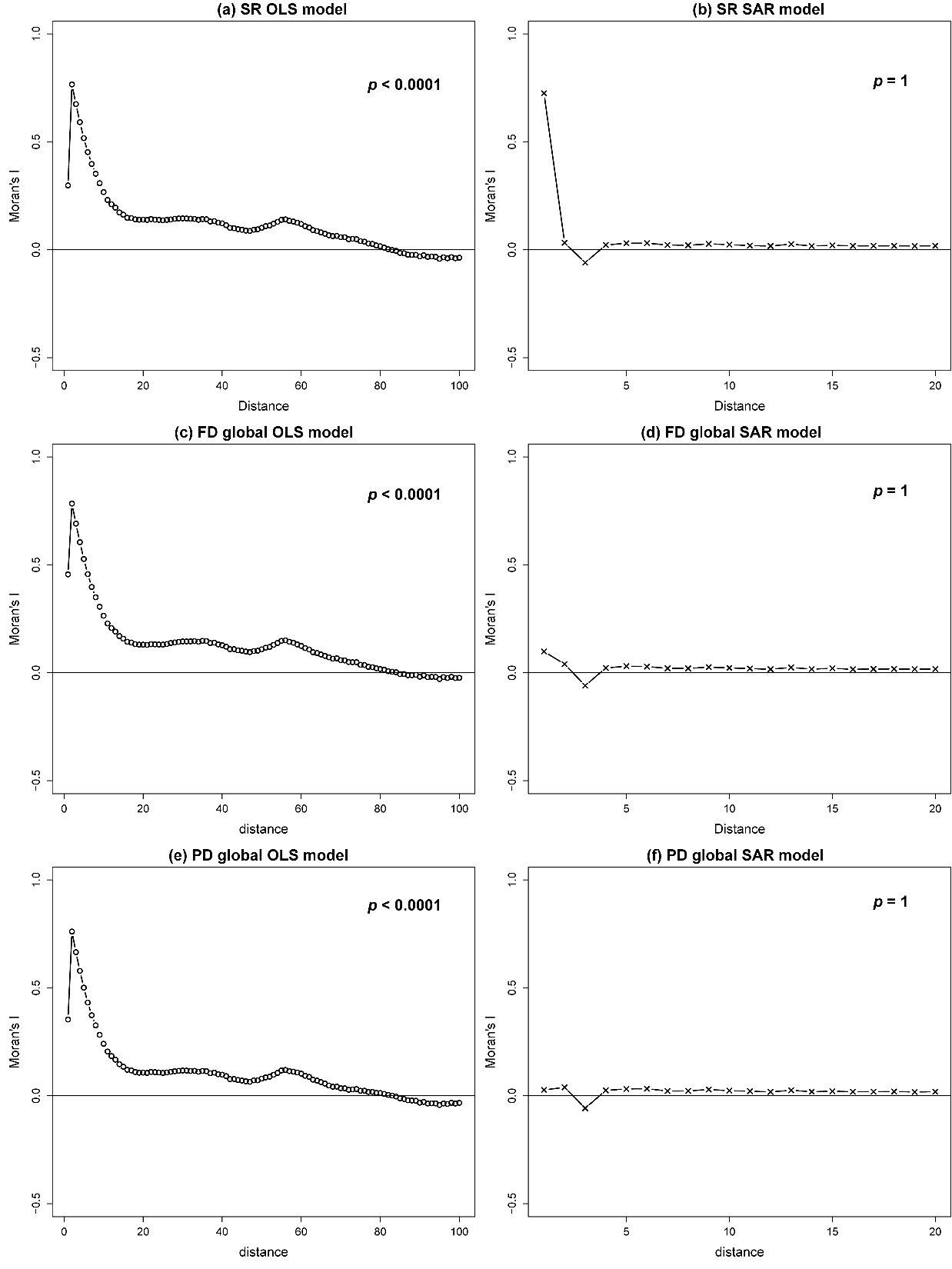

**Fig. S8** Correlograms of Maron’s *I* for the global species richness, functional and phylogenetic diversity ordinary least squares (OLS) and simultaneous autoregressive (SAR) models, as shown in Fig. 2. The summary of the models is shown in Table S2. The *p*-value in each subplot represents the significance of the Motan’s *I* test, and significant *p*-value indicates the residual of the model has strong spatial autocorrelation, and vice versa.

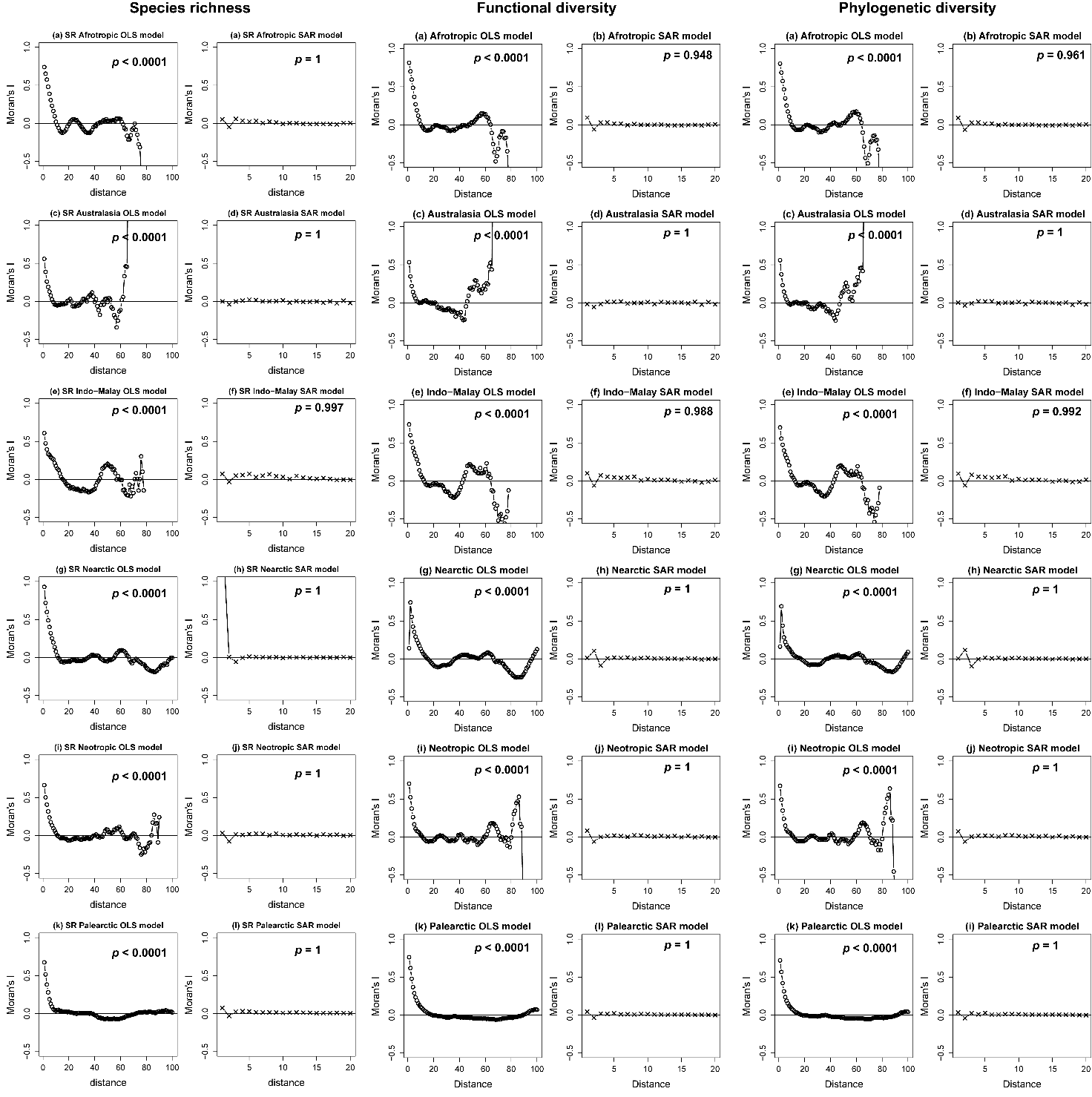

**Fig. S9** Correlograms of Maron’s *I* for the regional species richness, functional and phylogenetic diversity ordinary least squares (OLS) and simultaneous autoregressive (SAR) models (including anomaly variables and current mean annual temperature and annual precipitation), as shown in Fig. 3. The summary of the models is shown in Table S3. The *p*-value in each subplot represents the significance of the Motan’s *I* test, and significant *p*-value indicates the residual of the model has strong spatial autocorrelation, and vice versa.

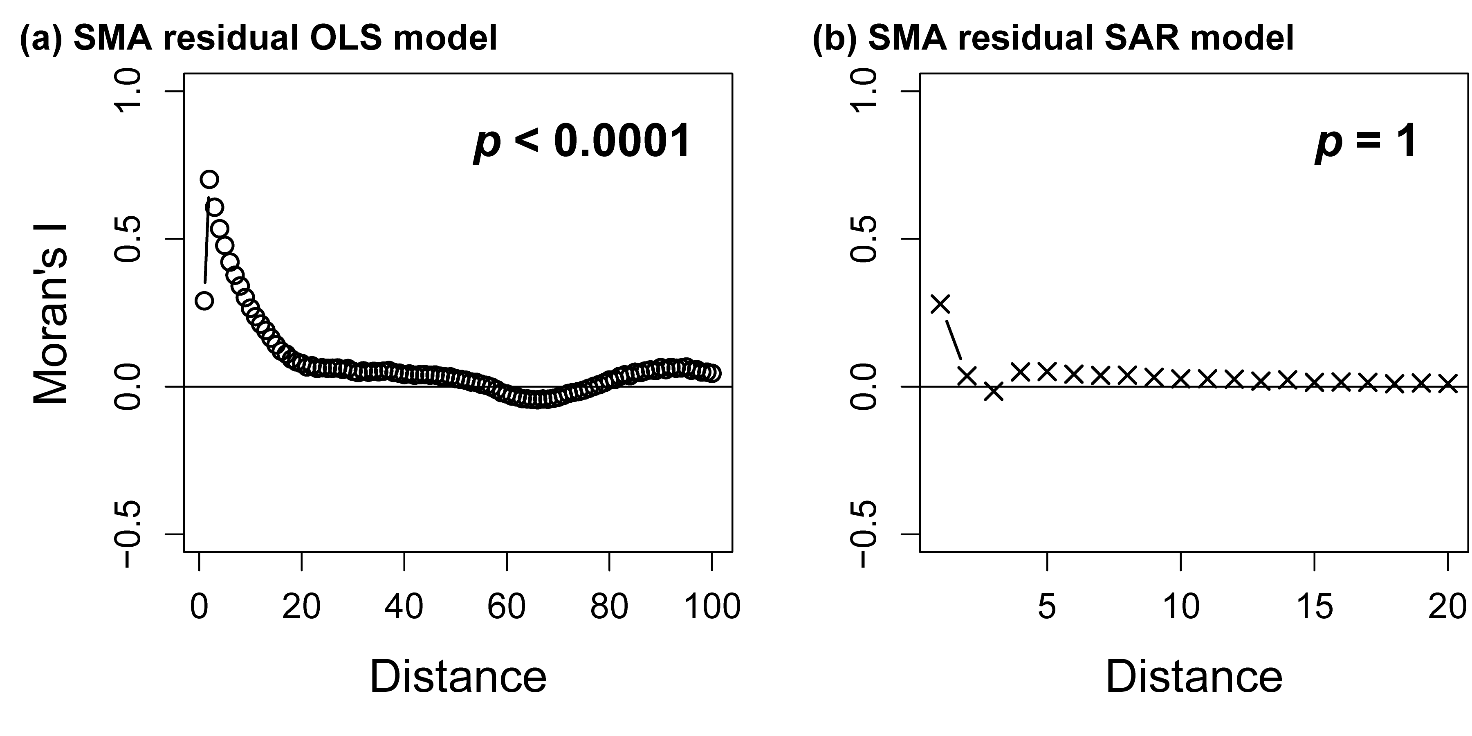

**Fig. S10** Correlograms of Maron’s *I* for global ordinary least squares (OLS) and simultaneous autoregressive (SAR) models of the residuals from the regression between functional and phylogenetic diversity (Fig. 3 & 4). The summary of the models is shown in Table S4. The *p*-value in each subplot represents the significance of the Motan’s *I* test, and significant *p*-value indicates the residual of the model has strong spatial autocorrelation, and vice versa.

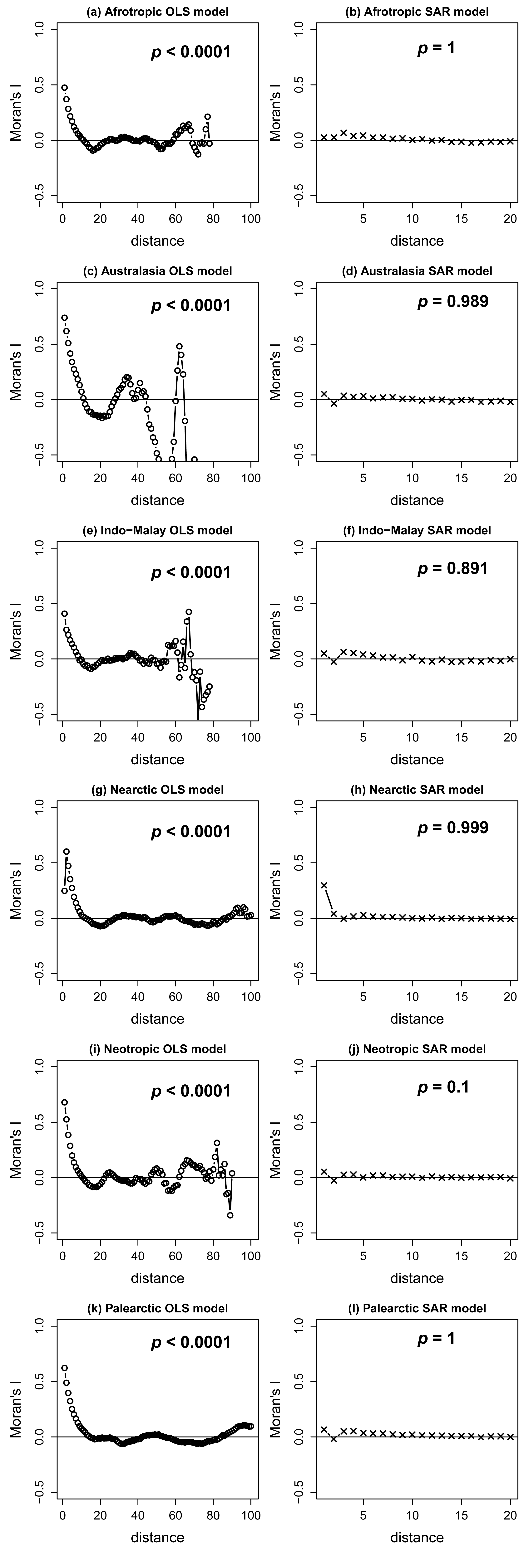
**Fig. S11** Correlograms of Maron’s *I* for the regional residual (from the regression between FD and PD) ordinary least squares (OLS) and simultaneous autoregressive (SAR) models, as shown in Fig. 4. The summary of the models is shown in Table S3. The *p*-value in each subplot represents the significance of the Motan’s *I* test, and significant *p*-value indicates the residual of the model has strong spatial autocorrelation, and vice versa.

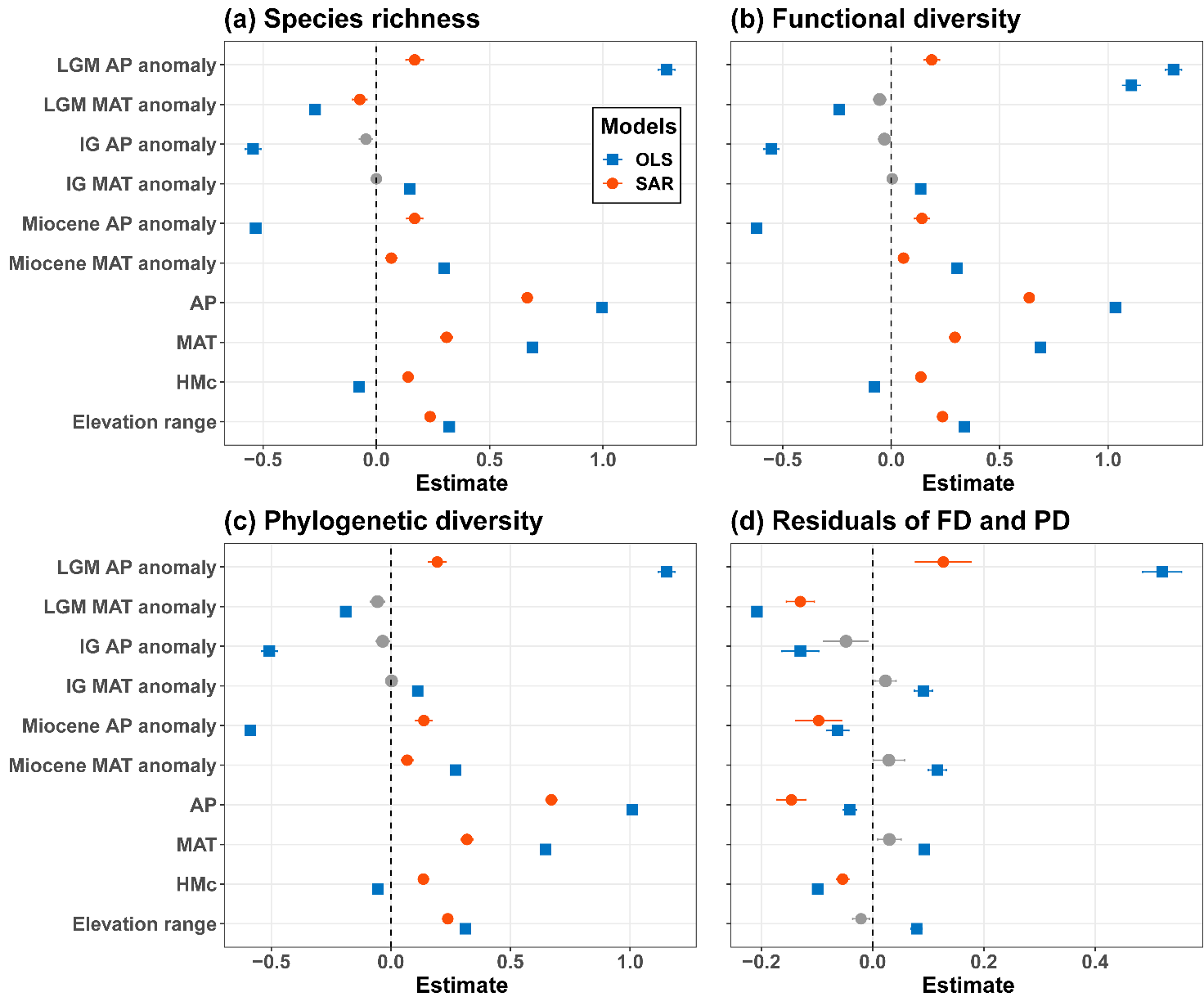

**Fig. S12** Effects of the tested environmental variables on tree (a) species richness (SR), (b) functional diversity (FD), (c) phylogenetic diversity (PD), and (d) residuals from the regression between FD and PD. Estimate (± 1standard error) of effects were obtained from global simultaneous autoregressive (SAR) models. Different colors and shapes indicate biogeographic regions. Non-significant variables (*p* > 0.05) are indicated in grey. Detailed results from both OLS and SAR models are shown in Tables S2 & S3. MAT: mean annual temperature; AP: Annual precipitation; HMc: human modification index; IG: Pleistocene Interglacial; LGM: Last Glacial Maximum.

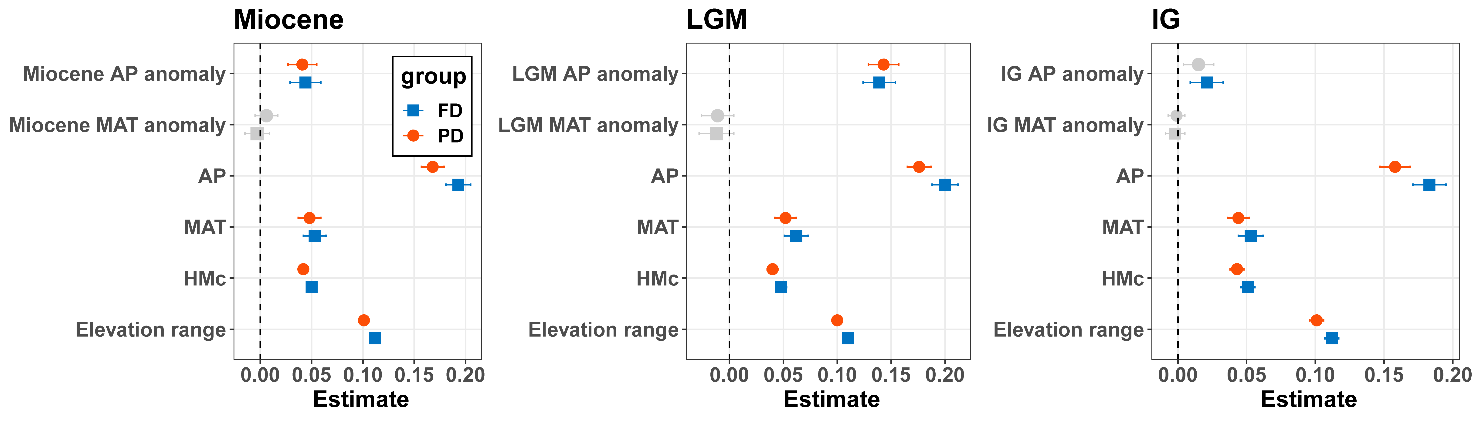

**Fig. S13** Effects of the each paleoclimate (both MAT and AP), contemporary MAT and AP, and non-climate variables on tree functional and phylogenetic diversity (FD and PD). Estimate (± 1standard error) of effects were obtained from simultaneous autoregressive (SAR) models. Non-significant variables (*p* > 0.05) are indicated in grey. Detailed results from OLS models are shown in Table S4. MAT: mean annual temperature; AP: Annual precipitation; HMc: human modification index; IG: Pleistocene Interglacial; LGM: Last Glacial Maximum.

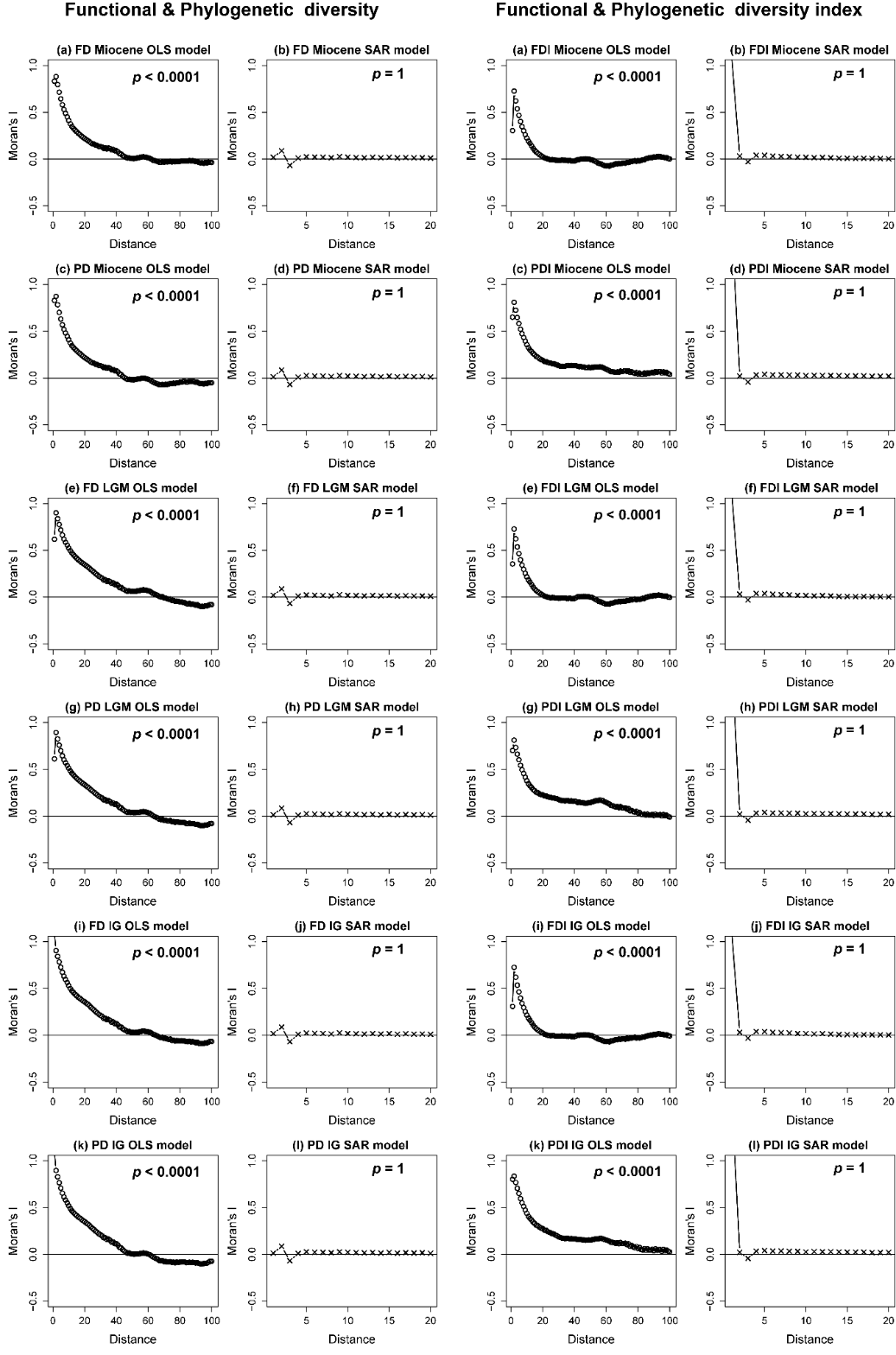

**Fig. S14** Correlograms of Maron’s *I* for the each paleoclimatic ordinary least squares (OLS) and simultaneous autoregressive (SAR) SES models, as shown in Fig. S12. The summary of the models is shown in Table S4. The summary of the models is shown in Table S4. The *p*-value in each subplot represents the significance of the Motan’s *I* test, and significant *p*-value indicates the residual of the model has strong spatial autocorrelation, and vice versa.

**Table S1** All environmental variables we considered. **Time** only applys to climatic variables and indicates when the variables are from. **Category** indicates the general climate situation in each time period or variable meanings (for the last three variables). Mya: Million years ago; kya: thousand years ago; MIS: Marine Isotope Stage.

| **Variables** | **Time** | **Category** | **Original resolution** | **Source** |
| --- | --- | --- | --- | --- |
| Annual mean temperature (^o^C) | Current climate (1970-2000) | Current | 30 arc-seconds | WorldClim.org |
| Annual Precipitation (mm) | Current climate (1970-2000) | Current |  | WorldClim.org |
| Annual mean temperature (^o^C) | Late Miocene climate (11.61-7.25 Mya) | Warm | 2.5 arc-minutes | Pound et al., 2011 |
| Annual precipitation (mm) | Late Miocene climate (11.61-7.25 Mya) | Warm |  | Pound et al., 2011 |
| Annual mean temperature (^o^C) | mid-Pliocene Warm period (3.264-3.025 Mya) | Warm | 2.5 arc-minutes | Hill 2015 & [paleoclim.org](http://paleoclim.org/) |
| Annual precipitation (mm) | mid-Pliocene Warm period (3.264-3.025 Mya) | Warm |  | Hill 2015 & [paleoclim.org](http://paleoclim.org/) |
| Annual mean temperature (^o^C) | Pliocene M2 (*ca*. 3.3 Mya) | Early cold | 2.5 arc-minutes | Dolan et al., 2015 & [paleoclim.org](http://paleoclim.org/) |
| Annual precipitation (mm) | Pliocene M2 (*ca*. 3.3 Mya) | Early cold |  | Dolan et al., 2015 & [paleoclim.org](http://paleoclim.org/) |
| Annual mean temperature (^o^C) | Pleistocene MIS19 (*ca*. 787 kya) | Interglacial | 2.5 arc-minutes | Brown et al., 2018 & [paleoclim.org](http://paleoclim.org/) |
| Annual precipitation (mm) | Pleistocene MIS19 (*ca*. 787 kya) | Interglacial |  | Brown et al., 2018 & [paleoclim.org](http://paleoclim.org/) |
| Annual mean temperature (^o^C) | Last Interglacial (*ca*. 130 kya) | Interglacial | 2.5 arc-minutes | Otto-Bliesner et al., 2006 & [paleoclim.org](http://paleoclim.org/) |
| Annual precipitation (mm) | Last Interglacial (*ca*. 130 kya) | Interglacial |  | Otto-Bliesner et al., 2006 & [paleoclim.org](http://paleoclim.org/) |
| Annual mean temperature (^o^C) | Last Glacial Maximum (*ca*. 21 kya) | Recent cold | 30 arc-seconds | Karger et al., 2017 & chelsa-climate.org |
| Annual precipitation (mm) | Last Glacial Maximum (*ca*. 21 kya) | Recent cold |  | Karger et al., 2017 & chelsa-climate.org |
| Global human modification index | - | Human disturbance | 30 arc-seconds | Kennedy et al., 2018 |
| Elevation range (m) | - | Topographic heterogeneity | 90 m | [<http://srtm.csi.cgiar.org/>](http://csi.cgiar.org/WhtIsCGIAR_CSI.asp) |
| Biogeographic regions | - | Evolutionary history |  | Morrone 2002 |

**Table** **S2** Summary of the global and biogeographic regional ordinary least squares (OLS) and simultaneous autoregressive (SAR) models for species richness, functional diversity and phylogenetic diversity, respectively (Fig. 1). HMc: human modification index; MAT: mean annual temperature; AP: Annual precipitation; IG: Pleistocene Interglacial. LGM: Last Glacial Maximum.

Excel file of “Table S2 Summary of OLS and SAR models of SR, FD and PD”

**Table S3** Summary of the global and biogeographic regional ordinary least squares (OLS) and simultaneous autoregressive (SAR) models for the residuals of the regression between functional diversity and phylogenetic diversity (Fig. 3). HMc: human modification index; MAT: mean annual temperature; AP: Annual precipitation; IG: Pleistocene Interglacial. LGM: Last Glacial Maximum.

Excel file of “Table S3 Summary of OLS and SAR models of residuals from regression between FD and PD”

**Table S4** Summary of the global ordinary least squares (OLS) and simultaneous autoregressive (SAR) models for the functional diversity and phylogenetic diversity in each paleoclimate scanerio (Fig. S12). HMc: human modification index; MAT: mean annual temperature; AP: Annual precipitation; IG: Pleistocene Interglacial. LGM: Last Glacial Maximum.

Excel file of “Table S4 Summary of OLS and SAR models of FD and PD from each paleoclimate”
